# Fam49b dampens TCR signal strength to regulate survival of positively selected thymocytes

**DOI:** 10.1101/2022.01.18.476719

**Authors:** Chan-Su Park, Jian Guan, Peter Rhee, Federico Gonzalez, Laurent Coscoy, Ellen A. Robey, Nilabh Shastri, Scheherazade Sadegh-Nasseri

## Abstract

The fate of developing T cells is determined by the strength of T cell receptor (TCR) signal they receive in the thymus. This process is finely regulated through tuning of positive and negative regulators in thymocytes. The Family with sequence similarity 49 member B (Fam49b) protein is a newly discovered negative regulator of TCR signaling that has been shown to suppress Rac-1 activity *in vitro* in cultured T cell lines. However, the contribution of Fam49b to thymic development of T cells is unknown. To investigate this important issue, we generated a novel mouse line deficient in Fam49b (Fam49b-KO). We observed that Fam49b-KO double positive (DP) thymocytes underwent excessive negative selection, whereas the positive selection stage was unaffected. This altered development process resulted in significant reductions in CD4 and CD8 single positive thymocytes as well as peripheral T cells. Interestingly, a large proportion of the TCRγδ^+^ and CD8αα^+^TCRαβ^+^ gut intraepithelial T lymphocytes were absent in Fam49b-KO mice. Our results demonstrate that Fam49b dampens thymocytes TCR signaling in order to escape negative selection during development, uncovering the function of Fam49b as a critical regulator of selection process to ensure normal thymocyte development.

## Introduction

Developing T cells in the thymus follow an ordered progression from CD4^-^CD8^-^ double negative (DN), to CD4^+^CD8^+^ double positive (DP), and finally to CD4 or CD8 single positive (SP) T cells [1]. Positive selection, negative selection, and CD4/CD8 lineage fate commitment of DP thymocytes rely on the strength of the interactions between TCR and self-peptides-MHC complexes [2]. Inadequate interactions lead to “death by neglect” whereas overly strong interactions lead to the elimination of thymocytes through “negative selection”. Thus, only those T cells receiving a moderate TCR signal strength are positively selected and further develop into mature T cells [2–4]. The TCR signal strength is also critical for CD4/CD8 lineage commitment. Enhancing TCR signaling in developing thymocytes favors development to the CD4 lineage, whereas reducing TCR signaling favors development of the CD8 lineage [5, 6].

While the majority of thymocytes bearing high affinity TCR for self-peptide MHC complexes undergo negative selection, not all self-reactive thymocytes follow this rule. Instead, these subsets of self-reactive non-deleting thymocytes are diverted to alternative T cell lineages through a process known as agonist selection [7, 8]. Several agonist selected T cell subsets has been defined including the CD8αα^+^TCRαβ^+^ intraepithelial lymphocytes (CD8αα^+^TCRαβ^+^ IELs), invariant natural killer T cells (iNKT cells), and Foxp3^+^ Regulatory T cells (Treg cells) [9–11]. Functionally, agonist selected T cells are thought to have a regulatory role in the immune system.

Actin cytoskeleton dynamics are important for multiple aspects of T cell function, including TCR signaling and adhesion, migration, differentiation, and execution of effector function [12–14]. In particular, actin cytoskeleton remodeling is required to provide scaffolding for TCR signaling proteins and for maintaining a stable immunological synapse between T cells and antigen presenting cells (APCs) [15–17]. However, the mechanisms that link actin cytoskeleton dynamics to the T cell signaling are not well understood. It has been reported that T cells cytoskeletal reorganization and regulation of actin dynamics at the immunological synapse are regulated by Rho family of small guanosine triphosphatases (Rho-GTPases) such as Rac [12]. Most members of Rho-GTPases exists in two conformational states between inactive (GDP-bound) and active (GTP-bound) [18]. The switch between the GDP- and GTP-bound is tightly regulated by guanine nucleotide exchange factors (GEFs) and GTPase-activating proteins (GAPs). GEFs activate Rho-GTPases by promoting the exchange of GDP for GTP, whereas GAPs inhibit Rho-GTPases by stimulating their GTP hydrolysis activity. Vav family proteins (Vav1, Vav2 and Vav3) are GEFs for Rac. Active Rac-1 transduce signals by binding to effector protein such as PAK and WAVE2 complex. Vav, Rac, and Pak play crucial roles in T cell development. For example, studies of mice lacking Vav-1 have shown that T cell development is partially blocked at pre-TCR β selection and is strongly blocked in both positive and negative selection [19–21]. Mice lacking both isoforms of Rac1 and Rac2 show defects in pre-TCR β-selection at DN thymocytes and positive selection of DP thymocytes [22, 23]. Mice lacking Pak2 show defects in pre-TCR β-selection of DN thymocytes, positive selection of DP thymocytes, and maturation of SP thymocytes [24].

Fam49b has been identified as an inhibitor of TCR signaling through binding with active Rac-1/2 in Fam49b-KO Jurkat T cells [25]. Those studies showed that lack of Fam49b led to hyperactivation of Jurkat T cells following TCR stimulation, as measured by the enhancement of CD69 induction, Rac-PAK axis signaling, and cytoskeleton reorganization [25]. Since TCR signaling strength controls thymocyte development, we hypothesized that Fam49b would be critical for thymocyte development *in vivo* and investigated this using a novel knockout mouse line. Here, we found the Fam49b was dispensable for positive selection but was required to prevent overly robust elimination of thymocytes at the negative selection stage, thus identifying Fam49b as a critical regulator of negative selection.

## Result

### Generation of Fam49b-KO mice and Fam49a-KO mice

To assess the role of Fam49b in T cell development, we generated Fam49b-KO mice by creating a premature stop codon in exon 6 of the *Fam49b* locus using CRISPR/Cas9 (**Fig. 1A**). Fam49a is a homologous protein that is ∼80% identical to Fam49b that has also been suggested to be involved in lymphopoiesis in zebrafish [26]. We generated Fam49a-KO mice in a similar manner by creating a stop codon in exon 7 of the *Fam49a* locus (**Fig.1B**). Immunoblot of spleen tissues confirmed that Fam49a or Fam49b expression was undetectable in Fam49a-KO mice or Fam49b-KO mice respectively in contrast to wild type (WT, C57BL/6J) mice (**Fig. 1C**). Real-time RT-PCR analysis of flow cytometry-sorted WT thymocytes subsets showed Fam49b is expressed broadly throughout thymic development, whereas Fam49a was mainly expressed in mature T cells (**Fig. 1D** and **Supplementary Fig. 1**). The expression of Fam49a was not detectable in WT thymocytes (**Fig. 1D**). Both Fam49a-KO and Fam49b-KO mice were fertile and did not show any apparent abnormalities.

**Figure 1.**
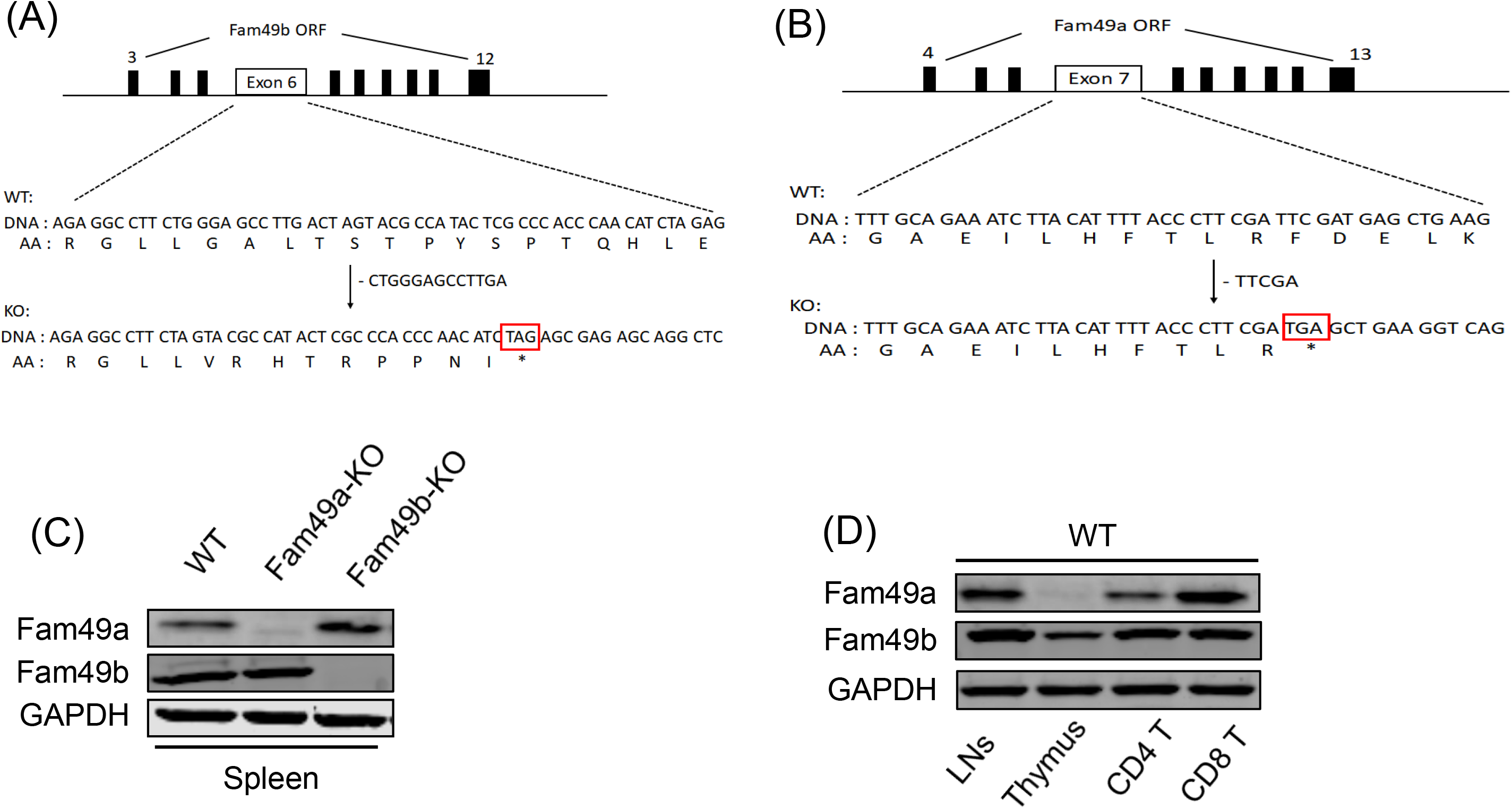
Generation of Fam49a-KO and Fam49b-KO mice with CRISPR/Cas9 and expression of Fam49a and Fam49b in mice. (**A**) Schematic diagram depicting the locations of guide RNAs (gRNAs) targeting the Fam49a. (**B**) Schematic diagram depicting the locations of guide RNAs (gRNAs) targeting the Fam49b. (**C**) Immunoblot analysis of Fam49a and Fam49b expression in spleen from WT, Fam49a-KO mice, and Fam49b-KO mice. The data are representative of three independent experiments. See also Figure 1 – source data 1. (**D**) Immunoblot analysis of Fam49a and Fam49b expression in lymph nodes, thymus, and peripheral CD4 T cells, and peripheral CD8 T cells from WT mice. The data are representative of three independent experiments. See also Figure 1 – source data 1.

### Defective T cell development in Fam49b-KO mice, but not Fam49a-KO mice

Flow cytometry analysis of cells isolated from lymph nodes showed that the frequency and number of peripheral CD4^+^ T cells and CD8^+^ T cells were significantly reduced in Fam49b-KO mice (**Fig. 2A**) compared to WT and Fam49a-KO mice. Notably, reduction in the number of CD8^+^ T cells was greater than that of CD4^+^ T cells. As a result. the ratio of CD4^+^ T cells over CD8^+^ T cells was increased in Fam49b-KO mice (**Fig. 2B**). In contrast, Fam49a-KO mice resembled WT mice in terms of T cell number and CD4/CD8 composition. Peripheral T cells in naïve mice can be divided into native and memory T subpopulation, which can be distinguished based on the expression of adhesion molecule CD62L and CD44. Assessment of the phenotype of peripheral T cells indicated that the reduction in T cell numbers was mainly due to a reduced number of naïve (CD44^lo^ CD62L^+^) CD4^+^ and CD8^+^ T cells in Fam49b-KO mice, while the size of the memory population was unchanged (**Fig. 2C**). Again, little difference was observed between Fam49a-KO and WT mice in terms of the phenotype of peripheral T cell subsets.

**Figure 2.**
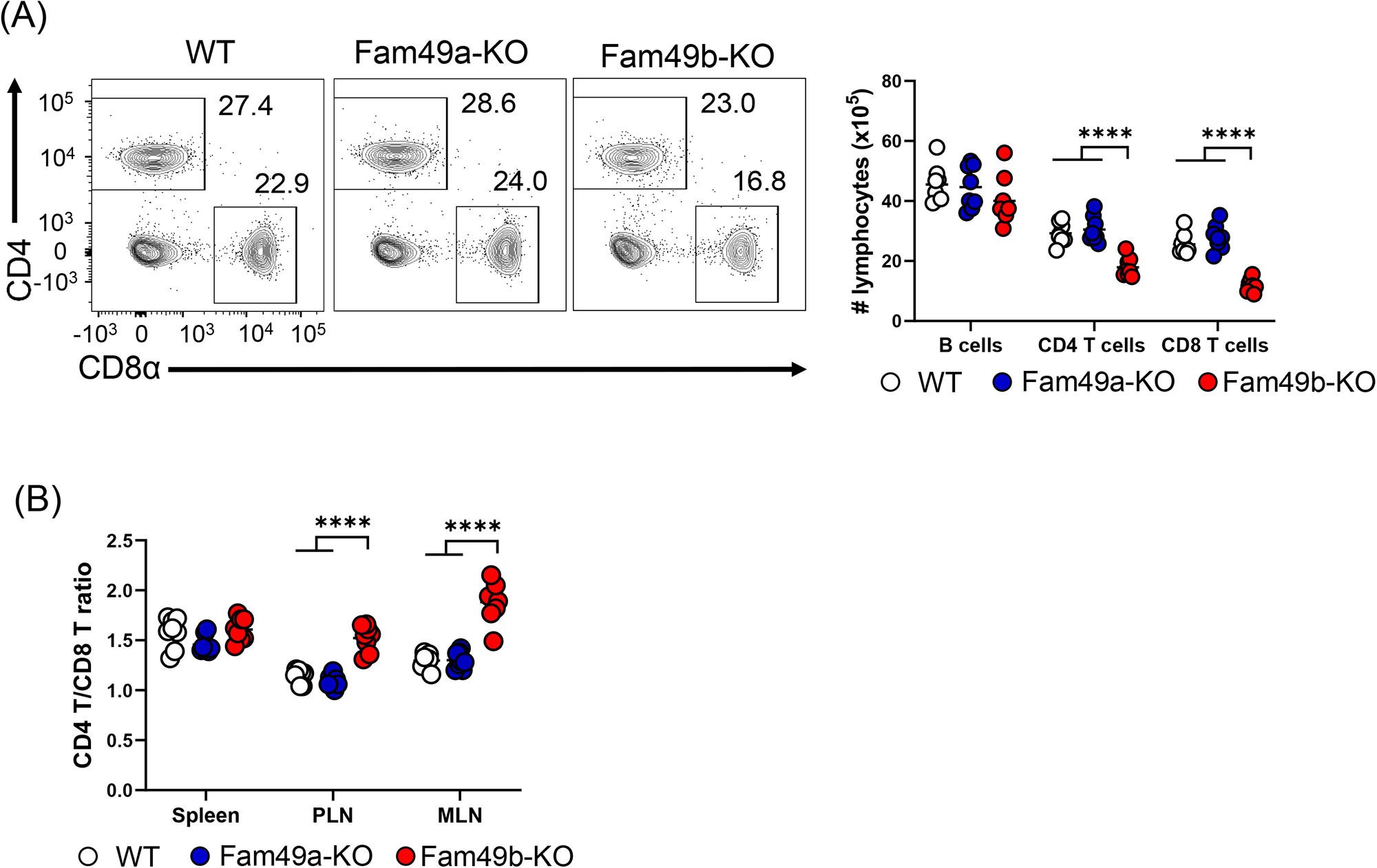
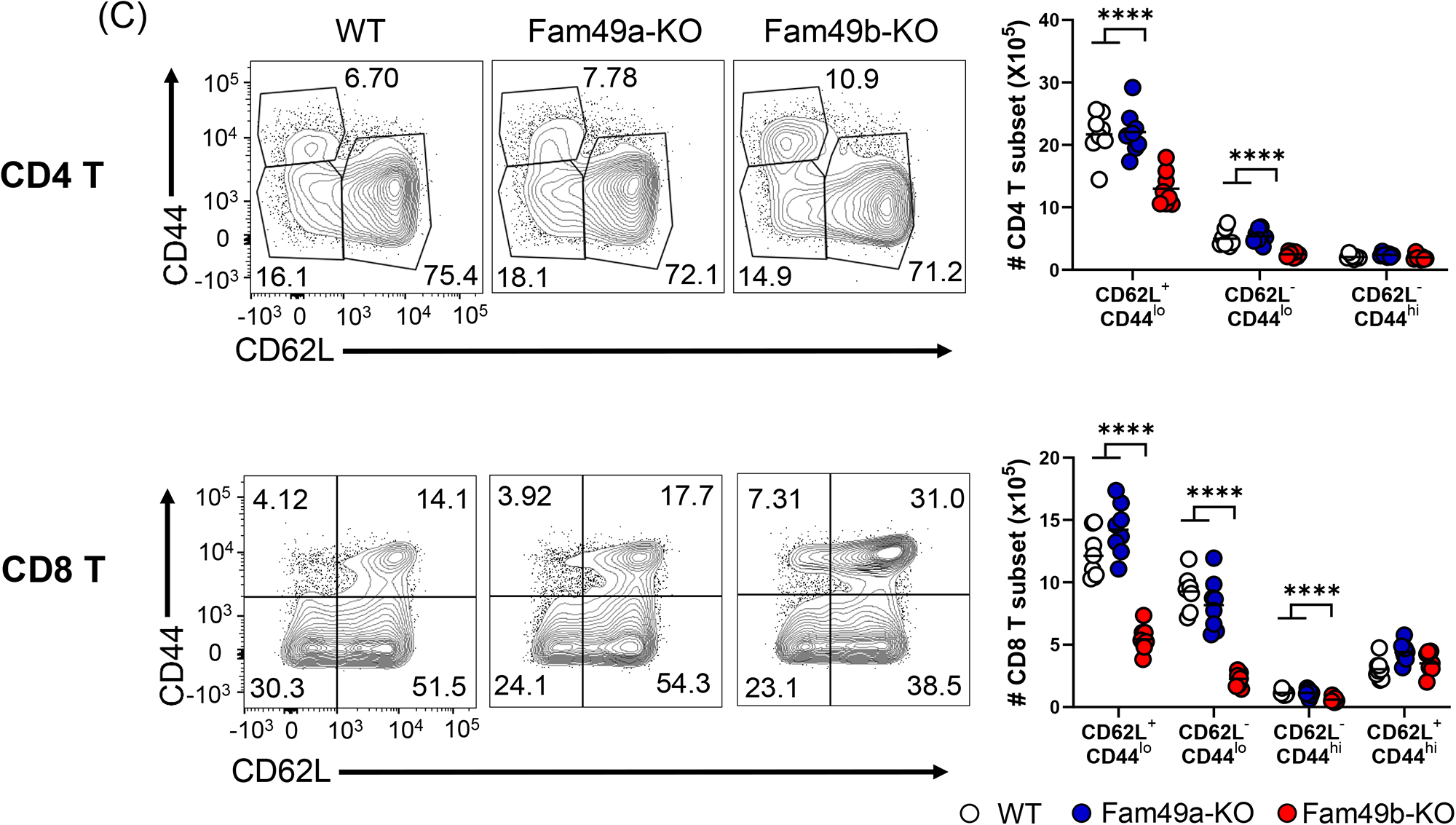
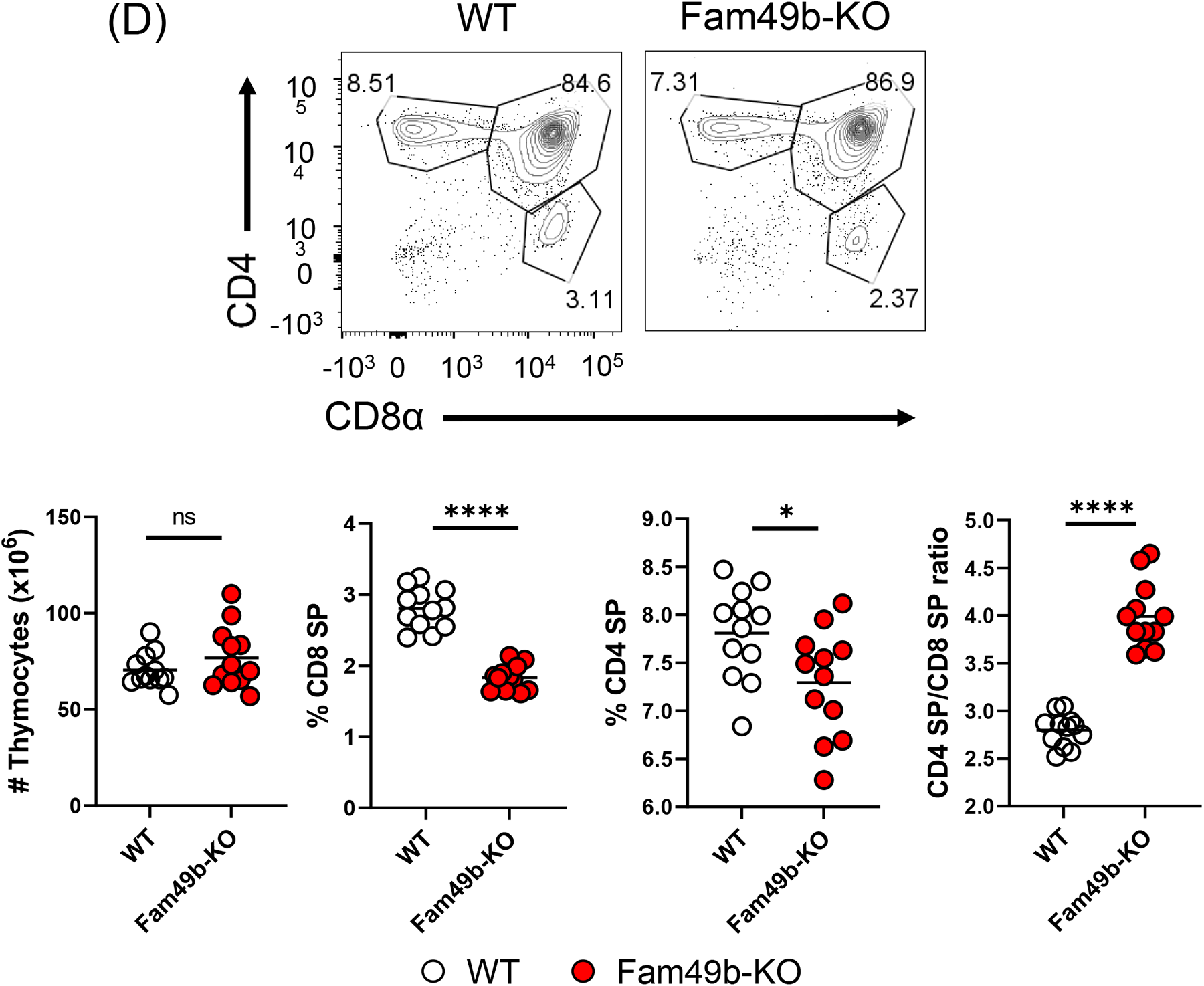
Reduced T cell numbers in Fam49b-KO mice, but not Fam49a-KO mice. (**A**) Flow cytometry profiles of the expression of CD4 and CD8 (**left**) and absolute number of lymphocytes in peripheral lymph nodes (**right**) from WT, Fam49a-KO, and Fam49b-KO mice. Numbers adjust to outlined areas indicate percentage of T cells among total lymphocytes. Each dot represents an individual mouse. Small horizontal lines indicate the mean of 8 mice. ****p<0.0001 (One-way ANOVA). Data are representative of four experiments. See also Figure 2 – source data 2. (**B**) Ratio of CD4 T cells over CD8 T cells in spleen, peripheral lymph nodes, and mesenteric lymph node in WT, Fam49a-KO, and Fam49b-KO mice. Each dot represents an individual mouse. Small horizontal lines indicate the mean of 8 mice. ****p<0.0001 (One-way ANOVA). Data are representative of four experiments. See also Figure 2 – source data 2. (**C**) Expression of CD44 and CD62L on T cells (**left**) and absolute number of T cell subset (**right**) in peripheral lymph nodes in CD4 T cells (**upper**) and CD8 T cells (**lower**) from WT, Fam49a-KO, and Fam49b-KO mice. CD4^+^ T subset with phenotype of naïve (CD62L^+^CD44^lo^), effector (CD62L^-^CD44^lo^), and memory (CD62L^-^ CD44^hi^) cells. CD8^+^ T subset with phenotype of naïve (CD62L^+^CD44^lo^), acute effector (CD62L^-^CD44^lo^), effector memory (CD62L^-^CD44^hi^), and central memory (CD62L^+^CD44^hi^). Numbers adjust to outlined areas indicate percentage of T cells subset among total T cells. Each dot represents an individual mouse. Small horizontal lines indicate the mean of 8 mice. ****p<0.0001 (One-way ANOVA). Data are representative of four experiments. See also Figure 2 – source data 2. (**D**) Flow cytometry analyzing the expression of CD4 and CD8 in thymocytes. Contour plots show percentage of CD8 SP and CD4 SP in total thymocytes (**upper**). Absolute number of total thymocytes (**lower left**), Percentage of CD4 SP and CD8 SP in total thymocytes (**lower middle**), and ratio of CD4 SP cells over CD8 SP cells was shown (**lower right**). Each dot represents an individual mouse. Small horizontal lines indicate the mean of 12 mice. *p=0.0295 and ****p<0.0001 (Mann-Whitney test). Data are representative of five experiments. See also Figure 2 – source data 2.

To further investigate if the decrease in naïve peripheral T cells subset in Fam49b-KO mice was due to defects of T cell development, we analyzed the surface expression of CD4 and CD8 on thymocytes. The frequencies of CD4 SP and CD8 SP cells were reduced and the ratios of CD4 SP to CD8 SP thymocytes were increased in Fam49b-KO mice thymi (**Fig. 2D** and **Supplementary Fig. 2**). These data indicate that Fam49b deficiency leads to impaired thymocyte development for both CD4^+^ and CD8^+^ T cells, with a more marked impact on the CD8^+^ T cell population. In contrast, loss of Fam49a showed little, if any, impacts on T cell numbers and cellularity in periphery, or T-cell thymic development. Given a lack of any phenotypic changes in Fam49a-KO mice T cells, together with an absence of Fam49a expression in thymus (**Fig. 1D**), we concluded that Fam49a is unlikely to play a significant role in T cell development. We therefore focused the remainder of our studies on the Fam49b-KO mice.

### Fam49b-KO thymocytes initiate positive selection but fail to complete development

Successful T cell development is a combined effort of both thymocytes and thymic microenvironment such as thymic epithelial cells and cytokine production. To determine if the effect of Fam49b deficiency on thymocytes development was thymocyte intrinsic or dependent on the extrinsic thymic microenvironment, we generated bone marrow chimera by injecting WT or Fam49b-KO CD45.2^+^ bone marrow cells into lethally irradiated WT CD45.1^+^ mice (B6.SJL-Ptprca Pepcb/BoyJ). A lower frequency of peripheral T cells (**Fig. 3A**) and increased ratio of peripheral CD4^+^ T over CD8^+^ T was observed in Fam49b-KO chimera mice compared to WT chimera mice (**Fig. 3B**). The Fam49b-KO thymocytes developed in WT thymic environment are like those developed in the germline Fam49b-KO environment in terms of both thymocyte and peripheral lymphocyte phenotypes. Therefore, the effect of Fam49b mutation on T cell development is predominantly due to thymocyte intrinsic functions.

**Figure 3.**
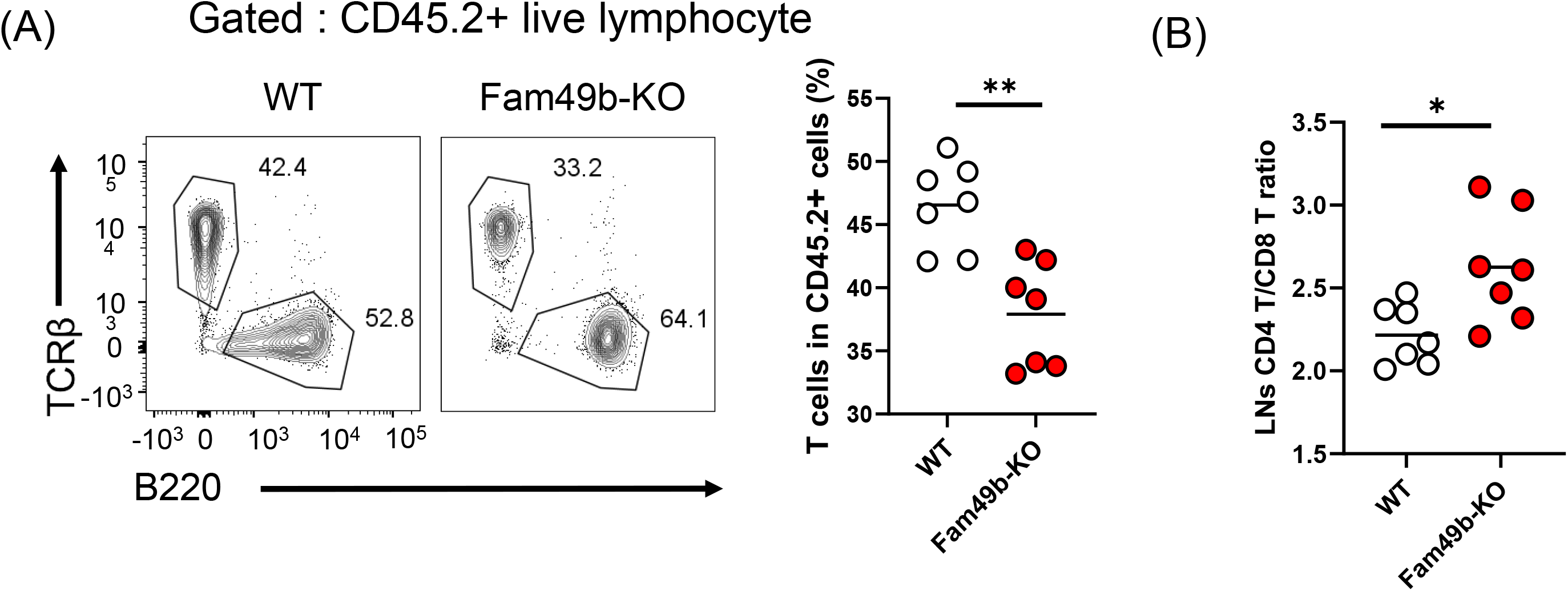
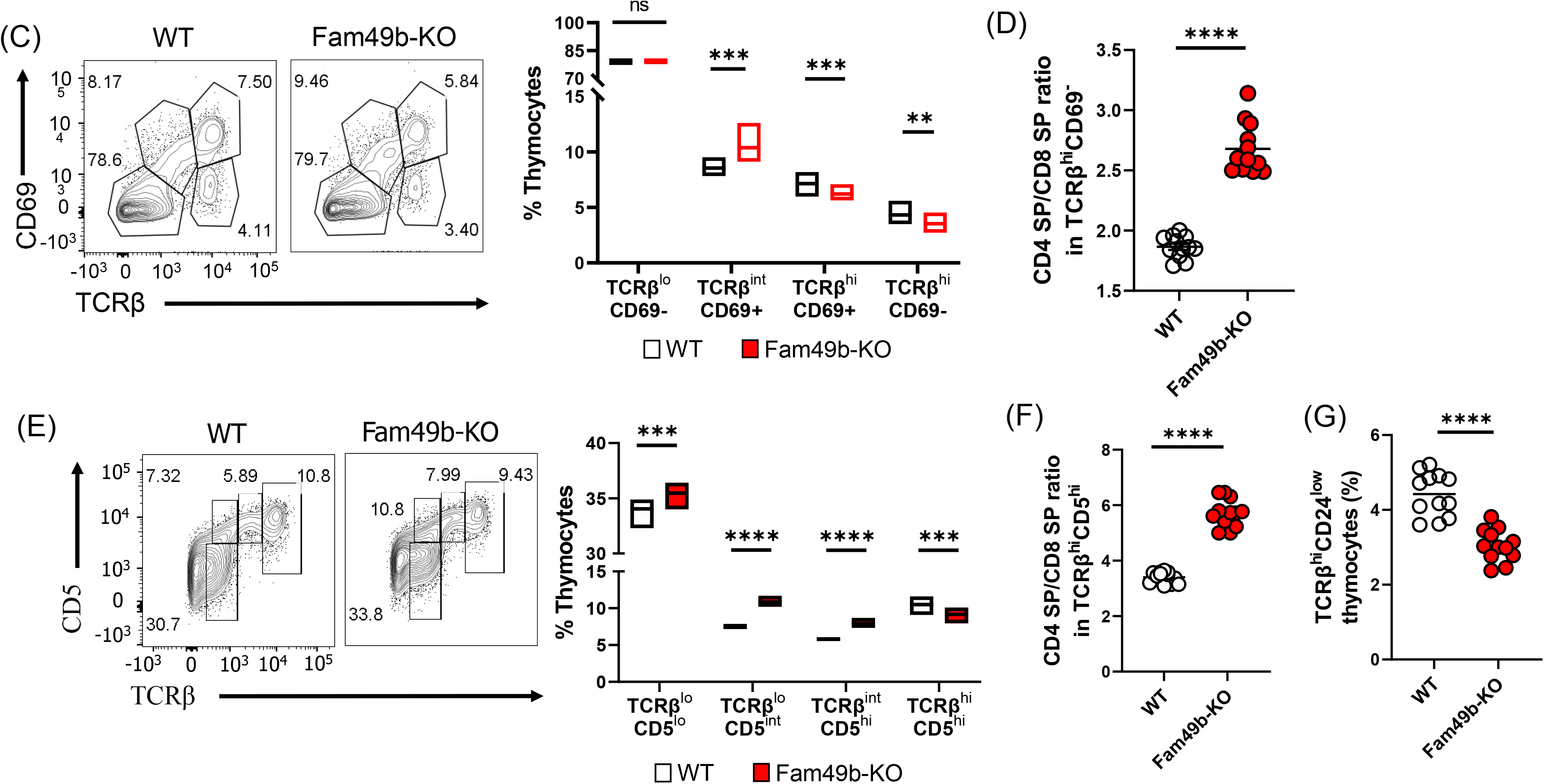
Defective thymic development in Fam49b-KO mice. (**A**) Expression of TCRβ and B220 expressing cells **(left**) and frequency of TCRβ expressing cells among CD45.2^+^ total lymph node cells from bone marrow chimera mice **(right**). Bone marrow from either WT or Fam49b-KO mice was injected *i.v*. into lethally irradiated CD45.1^+^ WT mice and chimeric mice were analyzed 8 weeks later. Small horizontal lines indicate the mean of 7 mice. **p=0.0047 (Mann-Whitney test). Data are pooled from two independent experiments. See also Figure 3 – source data 3. (**B**) Ratio of CD4 T cells over CD8 T cells in CD45.2^+^ total lymph node cells from bone marrow chimera mice. Bone marrow from either WT or Fam49b-KO mice was injected *i.v*. into lethally irradiated CD45.1^+^ WT mice and chimeric mice were analyzed 8 weeks later. Small horizontal lines indicate the mean of 7 mice. *p=0.0192 (Mann-Whitney test). Data are pooled from two independent experiments. See also Figure 3 – source data 3. (**C**) (**left**) Differential surface expression of CD69 and TCRβ was used to identify thymocyte population of different maturity in WT and Fam49b-KO mice. (**right**) Dot Plots show percentages of different thymocyte subpopulations in WT and Fam49b-KO mice. Numbers adjust to outlined areas indicate percentage of thymocytes subset among total thymocytes. Floating bars (min to max). horizontal lines indicate the mean of 12 mice. **p=0.0038 and ***p=0.0003 and ***p=0.0001 (Mann-Whitney test). Data are representative of five experiments. See also Figure 3 – source data 3. (**D**) Ratio of CD4 SP cells over CD8 SP cells in TCRβ^hi^CD69^-^ thymocyte subpopulation. horizontal lines indicate the mean of 12 mice. ****p<0.0001 (Mann-Whitney test). Data are representative of five experiments. See also Figure 3 – source data 3. (**E**) (**left**) Differential surface expression of CD5 and TCRβ was used to identify thymocyte population of different maturity in WT and Fam49b-KO mice. (**right**) Dot Plots show percentages of different thymocyte subpopulations from mice. Numbers adjust to outlined areas indicate percentage of thymocytes subset among total thymocytes. Floating bars (min to max). Horizontal lines indicate the mean of 12 mice. ***p=0.0005 and ***p=0.0002 and ****p<0.0001 (Mann-Whitney test). Data are representative of five experiments. See also Figure 3 – source data 3. (**F**) Ratio of CD4 SP cells over CD8 SP cells in TCRβ^hi^CD5^hi^ thymocyte subpopulation. Small horizontal lines indicate the mean of 12 mice. ****p<0.0001 (Mann-Whitney test). Data are representative of five experiments. See also Figure 3 – source data 3. (**G**) Frequency of TCRβ^hi^CD24^low^ thymocyte subpopulation among total live thymocytes. Small horizontal lines indicate the mean of 12 mice. ****p<0.0001 (Mann-Whitney test). Data are representative of five experiments. See also Figure 3 – source data 3.

Next, we sought to determine which step of T cell development was altered in Fam49b-KO mice. We thus subdivided thymocytes into four stages based on the differential expression of TCRβ and CD69 expression (**Fig. 3C** and **Supplementary Fig. 2**) [27]. The proportion of stage 1 thymocytes (TCRβ^lo^CD69^-^), which include the DN and pre-selection DP cells, was similar between WT and Fam49b-KO mice. The percentage of stage 2 thymocytes (TCRβ^int^CD69^+^), which represent transitional DP undergoing TCR-mediated positive selection, was significantly higher in the Fam49b-KO mice. The proportion of late stage thymocytes including the post-positive selection (TCRβ^hi^CD69^+^) and the mature thymocytes (TCRβ^hi^CD69^-^) was markedly decreased (**Fig 3C**). Consistent with our observation in periphery, the increased ratio of CD4 SP to CD8 SP was observed among the late stage thymocytes (TCRβ^hi^CD69^+^ and TCRβ^hi^CD69^-^) in Fam49b-KO mice (**Fig 3D**). These data show that the post-positive-selection process is impaired in Fam49b-KO thymocyte.

We further distinguished the pre-and post-positive selection populations by expression of cell surface TCRβ and CD5 (**Fig. 3E** and **Supplementary Fig. 3**) [27]. These markers define a developmental progression: stage 1 (TCRβ^lo^CD5^lo^) represents the pre-selection phase of DP thymocytes, and Stage 2 (TCRβ^lo^CD5^int^) are cells initiating positive selection. Stage 3 (TCRβ^int^CD5^hi^) represents thymocytes in the process of undergoing positive selection, and Stage 4 (TCRβ^hi^CD5^hi^) consists primarily of post-positive selection SP thymocytes. We observed that all the early phase populations (TCRβ^lo^CD5^lo^, TCRβ^lo^CD5^int^, TCRβ^int^CD5^hi^) increased significantly in proportion in Fam49b-KO, whereas the post-positive selection SP thymocytes (TCRβ^hi^CD5^hi^) were markedly decreased (**Fig. 3E**). Similarly, an increased ratio of CD4 SP to CD8 SP was observed in the post-positive selection population (TCRβ^hi^CD5^hi^) in Fam49b-KO thymocytes (**Fig. 3F**). This phenotype was further verified by the observation of lower percentage of mature SP CD24^lo^TCRβ^hi^ cells in Fam49b-KO mice compared with WT mice (**Fig. 3G**). Taken together, these results suggest that positive selection remains mostly unaffected by lack of Fam49b molecule, while Fam49b plays a more important role in the later stages of T cell development in the thymus.

### Enhanced negative selection in Fam49b-KO thymocytes

Based on our observation that loss of Fam49b led to decreased mature thymocyte populations, together with evidence that Fam49b can negatively regulate TCR signaling [25], we hypothesized that enhanced clonal deletion due to elevated TCR signaling strength would lead to the loss of positively selected thymocytes in Fam49b-KO mice. To test this hypothesis, we assessed cleavage of caspase 3, one of the key apoptosis events during clonal deletion (**Supplementary Fig. 4**) [28]. In the thymus, caspase 3 is cleaved in the apoptotic cells due to either clonal deletion (i.e. negative selection) or death by neglect (i.e. failed positive selection). To distinguish between these two fates, we stained the cells for TCRβ and CD5 molecules which are upregulated upon TCR stimulation. Thus, cleaved-caspase3^+^TCRβ^hi^CD5^hi^ cells represent thymocytes undergoing clonal deletion, whereas cleaved-caspase 3^+^TCRβ^-^CD5^-^ cells represent thymocytes undergoing death by neglect. We observed that the frequency of cells undergoing clonal deletion was increased among Fam49b-KO thymocytes, whereas the frequencies of cells to be eliminated through death by neglect were similar between Fam49b-KO and WT mice (**Fig. 4A**).

**Figure 4.**
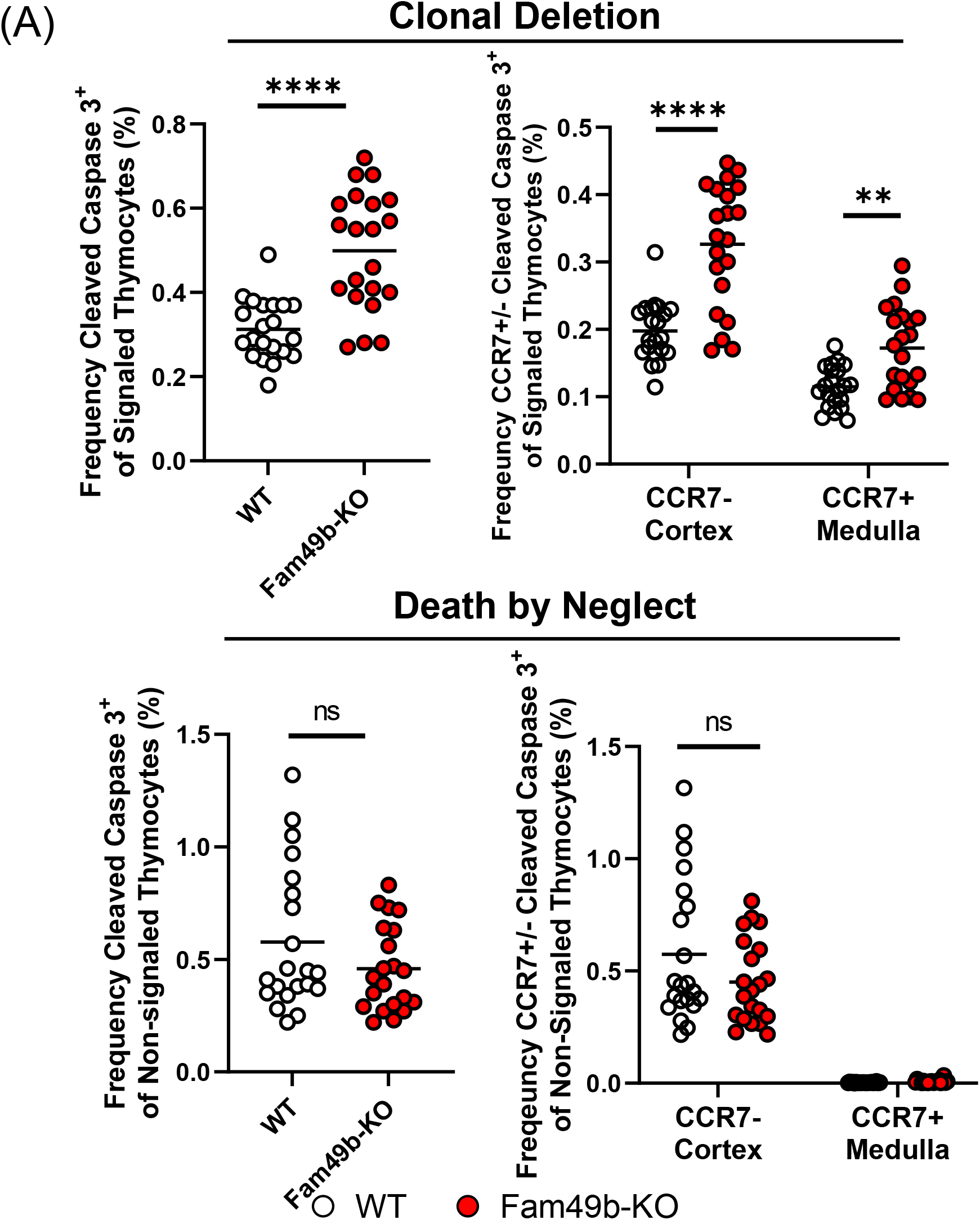
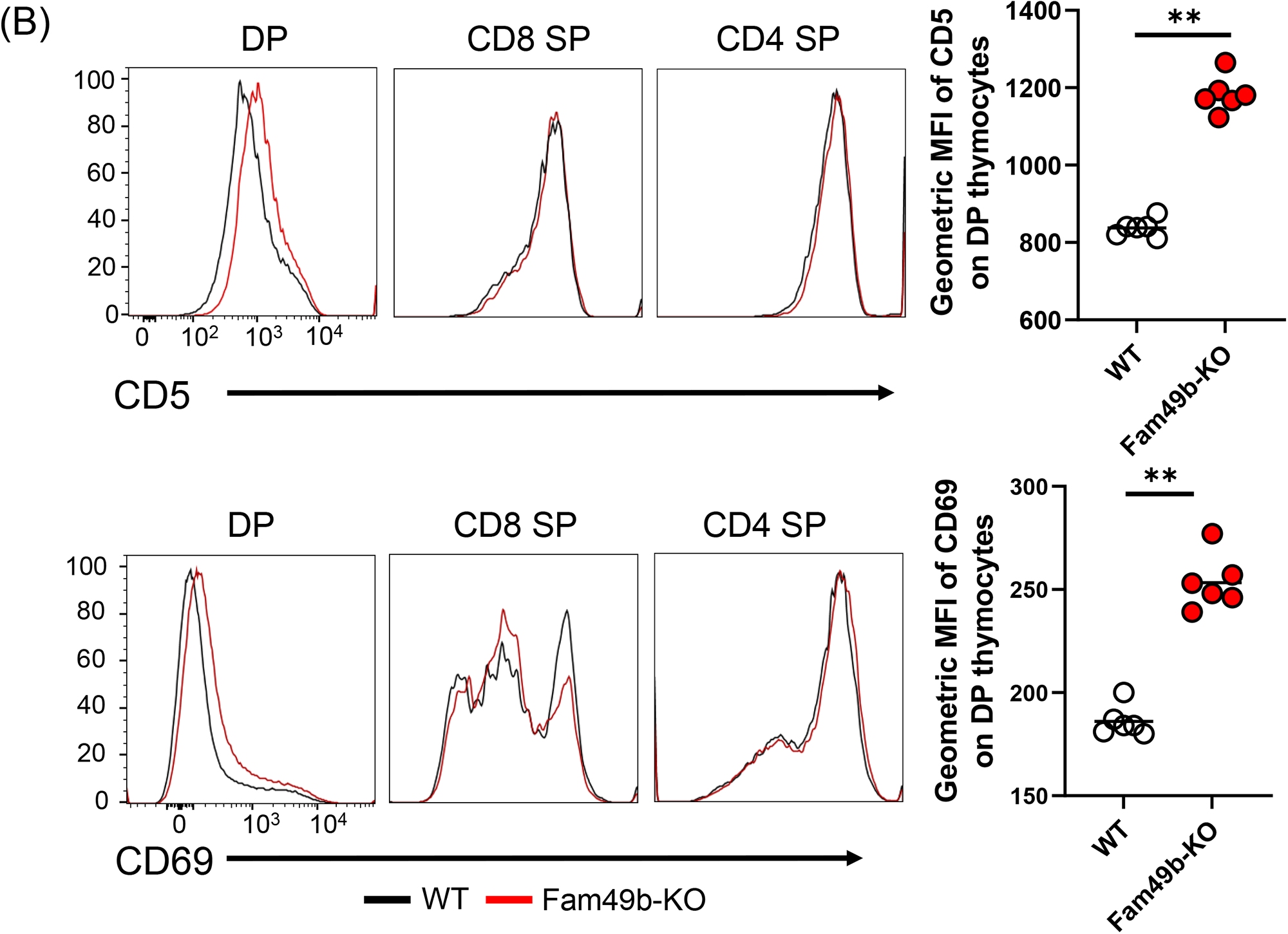
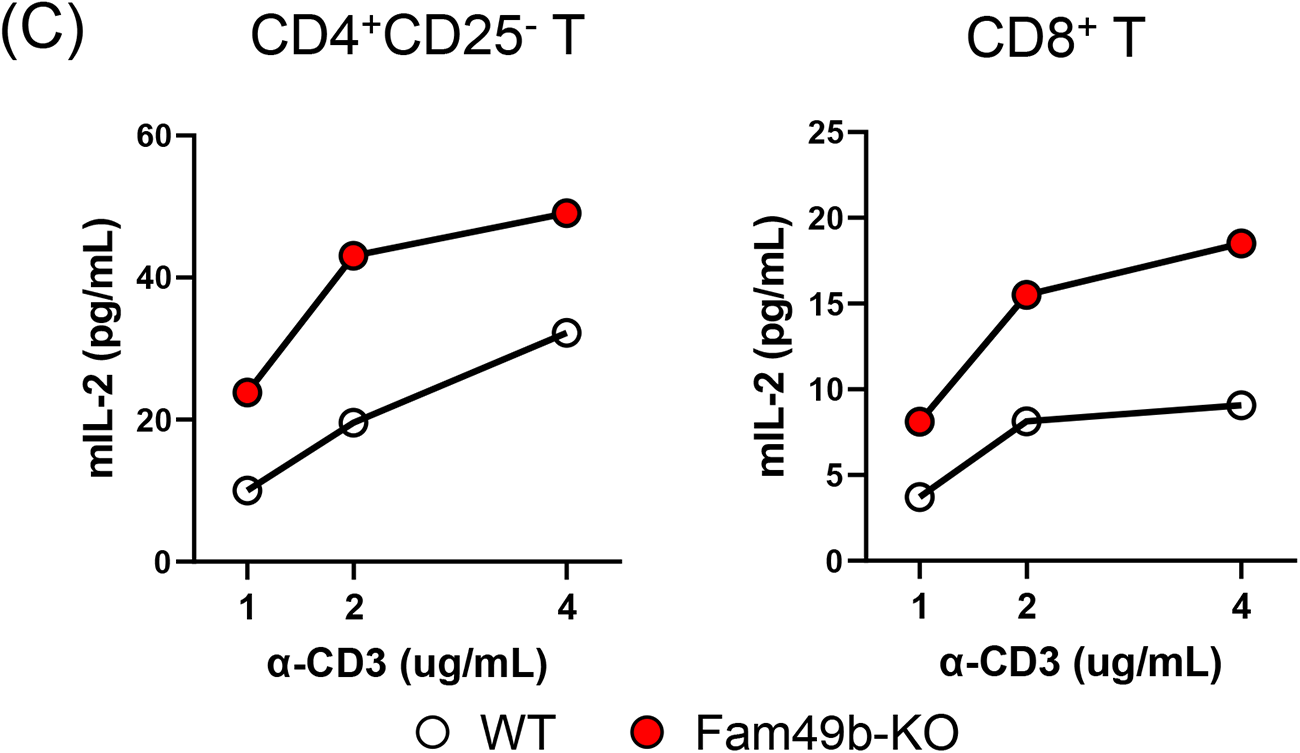
Enhanced negative selection due to elevated TCR signaling in Fam49b-KO thymocytes. (**A**) Frequency of cleaved caspase 3^+^ cells among TCRβ^hi^CD5^hi^ (Signaled, **upper left**) and TCRβ^-^CD5^-^ (Non-signaled, **lower left**) thymocytes. Frequency of CCR7^+^ cleaved caspase 3^+^ and CCR7^-^ cleaved caspase 3^+^ cells among TCRβ^hi^CD5^hi^ (Signaled, **upper right**) and TCRβ^-^CD5^-^ (Non-signaled, **lower right**) thymocytes. Small horizontal lines indicate the mean of 21 mice. **p=0.0017 and ****p<0.0001 (Mann-Whitney test). Data are pooled from three independent experiments. See also Figure 4 – source data 4. (**B**) Expression of activation marker CD5 on DP, CD4 SP, and CD8 SP thymocytes from WT and Fam49b-KO mice. (**upper**). Geometric MFI of CD5 on DP thymocytes (**upper right**). Expression of activation marker CD69 on DP, CD4 SP, and CD8 SP thymocytes from WT and Fam49b-KO mice (**lower**). Geometric MFI of CD69 on DP thymocytes (**lower right**). Small horizontal lines indicate the mean of 6 mice. **p=0.0022 (Mann-Whitney test). Data are representative of seven experiments. See also Figure 4 – source data 4. (**C**) Peripheral CD4^+^CD25^-^ T cells (**left**) or CD8^+^ T cells (**right**) were activated by immobilized anti-CD3ε (1, 2, 4 µg/mL) for 3 days, after which IL-2 in the supernatant was analyzed. Data are representative of four experiments. See also Figure 4 – source data 4.

Negative selection can occur in the thymic cortex as DP thymocytes are undergoing positive selection or in the thymic medulla after positive selection [29]. To determine whether loss of Fam49b led to increased deletion in the cortex or medulla, we stained the thymocytes for CCR7, which marks medullary thymocytes and is the receptor for the medullary chemokines CCL19/21 [28, 30]. The frequencies of cleaved-caspase3^+^CCR7^-^ cells and cleaved-caspase3^+^CCR7^+^ cells were significantly increased in the Fam49b-KO mice, suggesting that more thymocytes were eliminated through clonal deletion in both the cortex and medulla of Fam49b-KO thymus as compared with WT thymus (**Fig. 4A**).

Next, to determine if TCR-signal strength in Fam49b-KO thymocyte was increased, we assessed the surface expression of CD5 and CD69, two surrogate markers for TCR-signal strength [31, 32]. We found that both CD5 and CD69 expressions were upregulated on Fam49b-KO DP thymocytes, but not on CD4 SP and CD8 SP thymocytes (**Fig. 4B**), suggesting Fam49b-KO DP thymocytes had received stronger TCR signaling than the WT thymocytes. We next investigated the TCR-signaling strength of Fam49b-KO peripheral T cells by measuring IL-2 production in response to anti-CD3ε stimulation. Peripheral T cells were purified from spleen and lymph nodes from WT or Fam49b-KO mice and were stimulated in anti-CD3ε Ab coated plates for 3 days. We observed that IL-2 production was strongly elevated in Fam49b-KO CD4^+^CD25^-^ and CD8^+^ T cells compared to WT T cells (**Fig. 4C**). Despite the high IL-2 production of these cells, proliferation of both CD4^+^CD25^-^ and CD8^+^ Fam49b-KO T cells in response to anti-CD3ε stimulation *in vitro* were similar to that of WT T cells (**Supplementary Fig. 5**). In summary, enhanced TCR-signaling strength intrinsic to Fam49b-KO DP thymocytes leads to excessive clonal deletion in the cortex and medulla, resulting in the loss of naïve mature T cells in both thymus and periphery in the mice.

### Impaired development of natural IELs in Fam49b-KO mice

Some self-reactive thymocytes rely on strong TCR signaling to mature into unconventional T cell subsets through utilizing an alternative selection process known as agonist selection [33, 34]. Due to the robust effects of Fam49b deficiency on TCR-signaling strength, we investigated whether Fam49b affects the development of well-known agonist selected T cell subsets including CD8αα^+^TCRαβ^+^ IELs in small intestinal epithelium, iNKT cells in liver, and Treg cells in lymph nodes [9–11]. We found that all three T cell subsets were differentially affected by the loss of Fam49b. The percentage of CD8αα^+^TCRαβ^+^ IELs among IEL T cells was significantly decreased from 60% in WT mice to 30% in Fam49b-KO mice, whereas the frequency of liver iNKT cells was unaffected (**Fig. 5A**). The frequency of Treg among lymph node CD4^+^ T cells increased slightly from 16% to 20% in lymph nodes in Fam49b-KO mice, though the absolute number of Treg was ∼80% of the number in WT mice. Enhanced frequency of Treg seems to be a result of greater reduction of total CD4^+^ T cells compared to Treg (**Fig. 5A** and **Supplementary Fig. 6A**).

**Figure 5.**
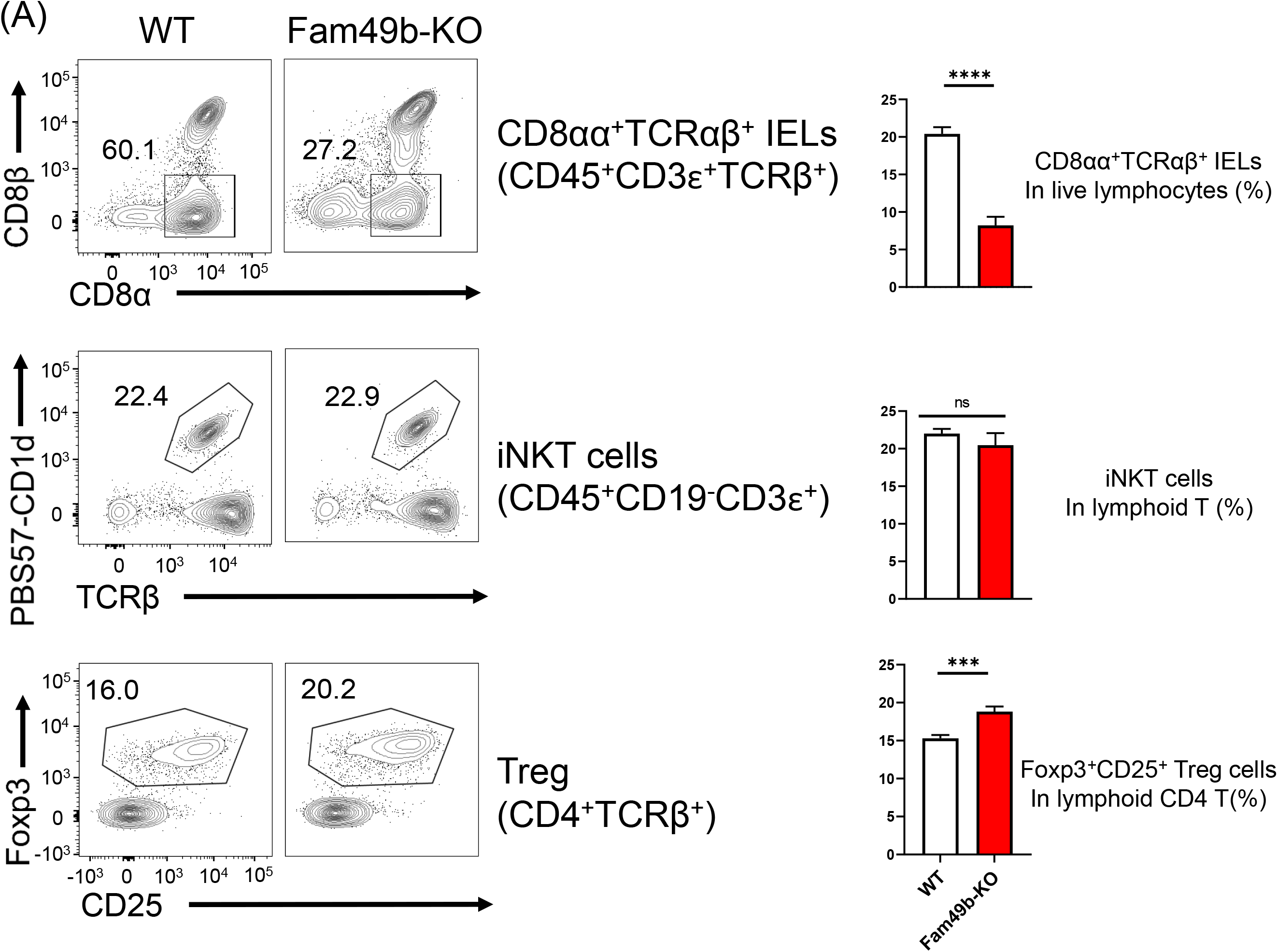
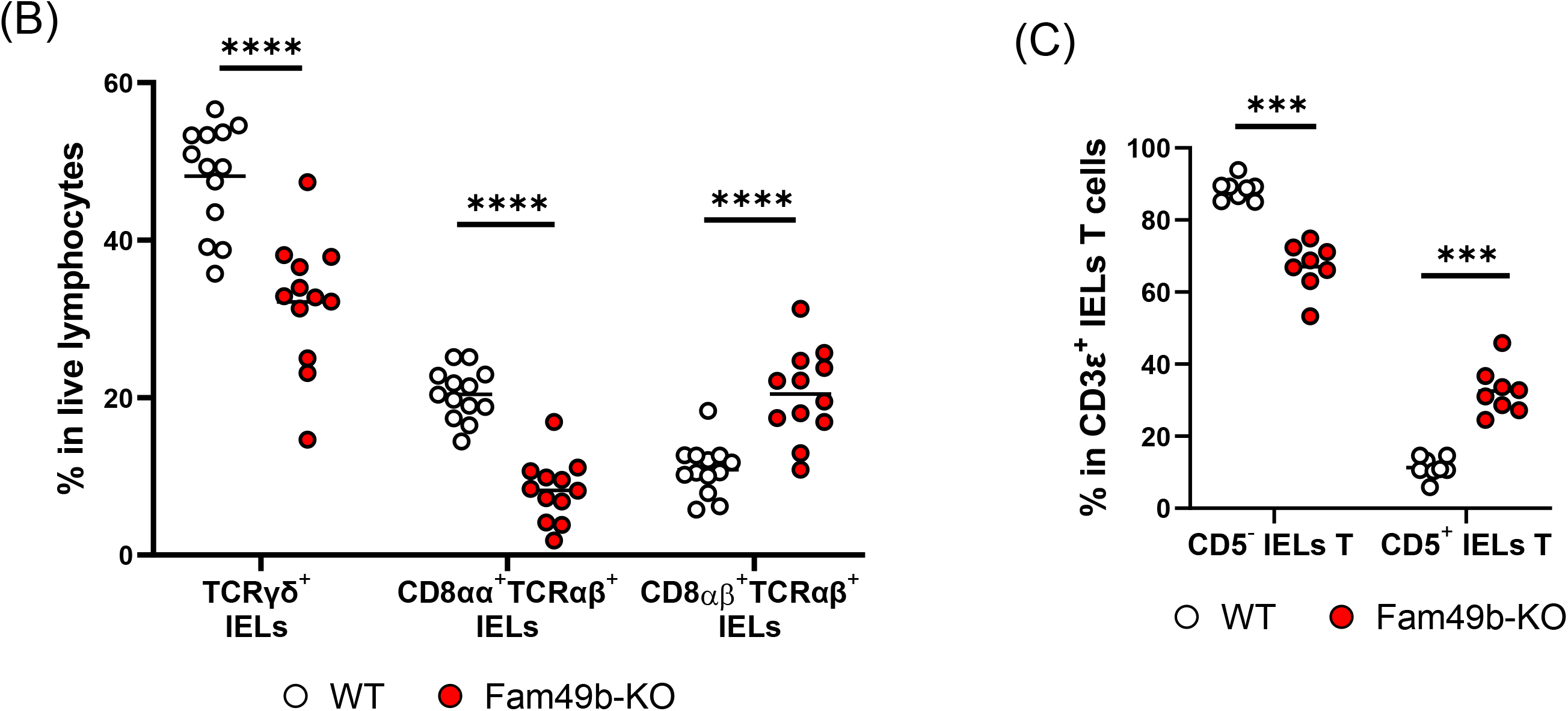
Fam49b-KO mice have lower frequency of CD8αα^+^TCRαβ^+^ and TCRγδ^+^ IELs T cells than WT mice. (**A**) Flow cytometry analysis of CD8αα^+^ TCRβ^+^ IELs T cells (**top**), CD1d-tetramer^+^ iNKT cells in the liver (**middle**), and Foxp3^+^CD25^+^ lymphoid regulatory T cells in the peripheral lymph nodes (**bottom**) from WT and Fam49b-KO mice. Right panels show average frequencies of each population among total lymphocytes or CD4 T cells. ***p=0.0003 and ****p<0.0001 (Mann-Whitney test). Data are pooled from seven independent experiments (CD8αα^+^ TCRβ^+^ IELs; mean and s.e.m, n=12∼13), representative of four experiments (iNKT cells; mean and s.e.m, n=6), or representative from seven independent experiments (Treg; mean and s.e.m, n=8). See also Figure 5 – source data 5. (**B**) Frequency of TCRγδ^+^ IELs T cells, CD8αα^+^TCRαβ^+^ IELs T cells, and CD8αβ^+^TCRβ^+^ IELs T cells among total live IELs cells in WT and Fam49b-KO mice. Each dot represents an individual mouse. Small horizontal lines indicate the mean of 12-13 mice. ****p<0.0001 (Mann-Whitney test). Data are pooled from seven independent experiments. See also Figure 5 – source data 5. (**C**) Frequency of CD5^+^ T cells and CD5^-^ T cells among total CD3ε^+^ IELs T cells in WT and Fam49b-KO mice. . Each dot represents an individual mouse. Small horizontal lines indicate the mean of 8 mice. ***p=0.0002 (Mann-Whitney test). Data are pooled from six independent experiments. See also Figure 5 – source data 5.

Gut IEL T lymphocytes are extremely heterogenous, and based on the differentiation mechanisms, can be subdivided into two major subpopulations including natural intraepithelial lymphocytes (natural IELs) and induced intraepithelial lymphocytes (induced IELs) [35]. Natural IELs home to gut immediately after thymic maturation. They are TCRγδ^+^ and TCRαβ^+^ T cells that can be either CD8αα^+^ or CD8αα^−^. In contrast, induced IELs arise from conventional peripheral CD8αβ^+^TCRαβ^+^ T cells and are activated post-thymically in response to peripheral antigens. The two populations can be distinguished by the expression of CD5; natural IELs are CD5^-^ and induced IELs CD5^+^. Based on our observation of the dramatic loss of CD8αα^+^TCRαβ^+^ IELs in Fam49b-KO mice, we postulated that other IEL subsets might be altered as well. Fam49b-KO mice showed a substantial reduction of natural IELs, including both the TCRγδ^+^ IELs as well as CD8αα^+^TCRαβ^+^ IELs (**Fig. 5B**), whereas the relative frequencies of induced IELs (CD8αβ^+^TCRαβ^+^ IELs) were increased (**Fig. 5C** and **Supplementary Fig. 6B**). These results suggest that Fam49b is involved in shaping the agonist-selected unconventional T cell populations and that Fam49b deficiency leads to substantial loss of the natural IELs, including CD8αβ^+^TCRαβ^+^ IELs and TCRγδ^+^ IELs.

## Discussion

Development of T cells is critically dependent on the strength of signaling through the TCR that lead to positive or negative selection [36, 37]. However, the roles of additional intracellular proteins and signaling pathways that regulate TCR signaling strength in the thymus have not been fully elucidated. Here, by studying the thymic development of T cells in Fam49b-KO mice, we report that Fam49b finetunes thymic selection by negatively regulating TCR signal-strength in the thymus and is essential for normal thymocyte development. Mice deficient in Fam49b developed severe T cell lymphopenia due to enhanced TCR-signaling in DP thymocytes. In Fam49b-KO thymus, post-positively selected population was significantly reduced, while generation of DN or immature DP thymocytes was mostly unaffected. We further confirmed that the loss of post-positive selection thymocytes in Fam49b-KO mice was due to enhanced clonal deletion instead of death by neglect. As a result, the frequencies of CD4 SP and CD8 SP cells in the Fam49b-KO thymi were significantly reduced.

While the medulla is a specialized site for negative selection, a substantial amount of negative selection occurs in the thymic cortex, overlapping in space and time with positive selection [38, 39]. We found that the frequency of thymocytes undergoing clonal deletion was significantly increased in Fam49b-KO thymus, while the frequency of thymocytes undergoing death by neglect remained the same. Moreover, most of thymocytes undergoing clonal deletion were CCR7^-^ cortex resident thymocytes (∼65%) in both WT and Fam49b-KO thymus. These data imply that Fam49b is needed immediately after the initial positive selection stage to serve as a ‘brake’ which dampens TCR signaling, thus helping to avoid negative selection. This ‘brake’, once taken out of the picture, leads to overexuberant clonal deletion and subsequent loss of a large proportion of the mature T cells.

Fam49b-KO DP thymocytes received stronger TCR signal compared to WT DP thymocytes. At the molecular level, Fam49b directly interacts with active Rac and negatively regulates its activity [25, 40, 41]. Rac plays key roles in cytoskeleton remodeling, signal transduction, and regulation of gene expression in thymocytes and peripheral T cells [42, 43]. The modulation of Rac activity by switching between its two conformational states, i.e., inactive (GDP-bound) and active (GTP-bound), is essential for multiple stages of thymocyte development and maturation. Previous studies suggested that Rac activity is important for β selection at DN thymocytes as well as positive and negative selection at DP thymocytes [22, 42, 44]. Moreover, transgenic mice that express constitutively active Rac-1 mutant revealed that Rac-1 activity could reverse the fate of thymocytes from positive to negative selection in the thymus [45]. Taken together, the phenotype similarities between active Rac-1 transgenic and our Fam49b-KO mice, and the association between Rac and Fam49b molecule, suggests that the impaired T-cell development in Fam49b-KO mice is likely a result of enhanced Rac activity in DP thymocytes.

How might enhanced Rac activity lead to the defective T-cell development in Fam49b-KO mice? Rac is known to regulate actin reorganization in T cells through binding with the Rac downstream effectors, such as PAK and WAVE2 complex [13]. The Pak2-deficient CD4 thymocytes showed weakened TCR-signaling strength as indicated by reduction of Nur77 expression in response to αCD3-stimulation [24], suggesting that the Rac-driven cytoskeleton remodeling is important for downstream events of TCR signaling. Negatively regulated Rac-driven cytoskeleton remodeling could attenuate protrusion and migration process in T cells. Fam49b-deficient cells showed increased cellular spread and reduced protrusion-retraction dynamics [40, 41]. Moreover, negative selection occurs via lengthy interactions between T cells and APCs, whereas positive selection are transient interactions [46]. Therefore, it is possible that altered cytoskeleton remodeling activity in Fam49b-KO thymocytes contributed to their elevated TCR-signaling strength and enhanced negative selection, perhaps by prolonging interactions with thymic APCs.

Among all the unusual phenotypes of peripheral T cells in Fam49b-KO mice, one surprising yet interesting observation was the significant loss of CD8αα^+^TCRαβ^+^ and TCRγδ^+^ IELs T cells. These T cell subsets were previously defined as unconventional T cells derived from self-reactive thymocytes that mature through agonist selection. The development of agonist-selected T cells relies on relatively strong and sustained TCR signaling which correlates with the magnitude of store-operated Ca2^+^ entry and NFAT activity [8, 33]. Yet it remains unclear why these cells that receive unusually high TCR signal are not eliminated through negative selection, but instead traffic into the gut and become IEL T cells [47, 48]. Interestingly, thymocytes undergoing agonist selection into CD8αα^+^ TCRαβ^+^ IELs T cells exhibited a rapid and confined migration pattern, in contrast to negatively selecting cells, which showed arrested migration [49]. It is tempting to speculate that overactivation of Rac-1 in Fam49b-KO mice might lead to negative selection of IEL precursors, perhaps by favoring migratory arrest over confined migration after encountering with agonist ligands.

In conclusion, Fam49b is critical for the thymic development of conventional T cells as well as unconventional natural IELs T cells. Interestingly, the function of Fam49b is restrained to the late-phase T cell development, where it dampens TCR signals to avert negative selection of DP thymocytes. The action of Fam49b is key in distinguishing positive from negative selection in thymic development. Our study offered insights on the association between modulation of TCR-signaling strength, cytoskeleton remodeling, and thymic development process.

## Materials and Methods

### Mice

C57BL/6 (WT) and CD45.1 mice (B6.SJL-Ptprca Pepcb/BoyJ, stock no: 002014) were purchased from the Jackson Laboratory and bred in house. Fam49a-KO and Fam49b-KO mice were generated by CRISPR/Cas9 gene-editing technology. The construct was electroporated into embryonic stem cells at the University of California at Berkeley gene targeting facility. All mouse procedures were approved by the Johns Hopkins University Animal Care and Use Committee and were following relevant ethical regulations.

### Antibodies and reagents

– Western blotting: anti-Fam49a (1103179) Millipore Sigma (St. Louis, MO); anti-Fam49b (D-8) Santa Cruz (Dallas, Texas); anti-GAPDH (ab9485) Abcam (Waltham, MA); anti-mouse IgG (926-68072), anti-rabbit IgG (926-32211) Li-cor (Lincoln, NE).
– Stimulation: anti-CD3e (145-2C11) was from BD Biosciences (San Jose, CA).
– Flow cytometry : anti-CD3e (17A2), anti-CD4 (RM4-5), anti-CD5 (53-7.3), anti-CD8α (53-6.7), anti-CD8β (53-5.8), anti-CD19 (ID3), anti-CD24 (M1/69), anti-CD25 (PC61), anti-CD44 (IM7), anti-CD45 (30-F11), anti-CD45.1 (A20), anti-CD45.2 (104), anti-CD45R/B220 (RA3-6B2), anti-CD62L (MEL-14), anti-CD69 (H1.2F3), anti-CD197/CCR7 (4B12), anti-Ly6G (IA8), anti-Ly-6C (HK1.4), anti-NK1.1 (PK136), anti-TCRβ (H57-597), anti-TCRγδ (GL3), anti-H-2Kb (AF6-88.5); anti-CD16/32 (93) BioLegend (San Diego, CA); anti-CD11b (M1/70), anti-T-bet (4B10) BD Bioscience ; anti-Cleaved Caspase 3 (D3E9) Cell signaling (Danvers, MA); Foxp3 (FJK-16s) eBioscience (San Diego, CA).
– Tetramerization: PBS-57 loaded mouse CD1d monomers were synthesized by the Tetramer Core Facility of the US National Institute of Health, Streptavidin-APC (PJ27S) and Strepavidin-RPE (PJRS27) were purchased from Prozyme (Agilent, Santa Clara, CA).

### Immunoblotting

Cell extracts were prepared by resuspending cells in PBS, then lysing them in RIPA buffer containing protease inhibitor cocktail (ThermoFisher Scientific, Waltham, MA). Protein concentrations were determined with the BCA Protein Reagent Kit (Pierce, ThermoFisher Scientific), after which 2-mercaptoethanol and 4x Laemmli Sample buffer (Bio Rad, Hercules, CA) were added and the samples were boiled. Western blotting was performed according to standard protocols using anti-Fam49a pAb, and anti-Fam49b mAb, and anti-GAPDH pAb. IRDye800CW conjugated goat anti-rabbit and IRDye680RD conjugated donkey anti-mouse were used as secondary antibodies. The membrane was scanned with the Odyssey Infrared Imaging System (Li-cor, model 9120)

### Real-time RT-PCR

The subsets of C57BL/6 thymocytes was collected using BD FACSAria II Cell sorter by the Ross Flow Cytometry Core Facility of the johns Hopkins. DN1 cells were gated as CD25^−^CD44^hi^; DN2 cells were gated as CD25^+^CD44^int–hi^; DN3 cells were gated as CD25^+^CD44^neg–lo^; and DN4 cells were gated as CD25^−^CD44^−^. Total RNA was isolated using the RNeasy Plus Micro Kit (Qiagen, Germantown, MD) and cDNA was amplified by SuperScript IV First Strand Synthesis (Invitrogen, ThermoFisher Scientific) according to the manufacturer’s instructions. Real-time PCR was performed using SYBRgreen PCR Master Mix (Applied Biosystems, ThermoFisher Scientific) and the ViiA 7 Real-Time PCR System (Applied Biosystems, ThermoFisher Scientific). Fam49b primers were forward, 5’-AGGAGCTGGCCACGAAATAC-3’, and reverse, 5’-GGCGTACTAGTCAAGGCTCC-3’. Results were normalized to β-actin expression with the 2−ΔCt method.

### Isolation of immune cells

Small-intestine IELs were isolated as previously described [50] : Changes were made as follow: DTT (BP172-5, Fisher Scientific) was used instead of DTE. Immune cells were collected from the interface of the 44% and 67% Percoll gradient and characterized by flow cytometry. Hepatic lymphocytes were isolated using Liver dissociation kit (Miltenyi Biotec, Auburn, CA) according to the manufacturer’s instructions. Samples were resuspended in 33% Percoll and spun, and the cell pellet was collected and labeled for flow cytometry.

### IL-2 ELISA and proliferation assay

T cells were purified from spleen and lymph nodes by negative selection using Miltenyi Biotech MACS cell isolation kit. Cells were labeled with 5 µM CellTrace Violet Cell Proliferation Kit (Invitrogen, ThermoFisher Scientific) at room temperature for 10 mins. Staining cells was washed and cultured anti-mouse CD3ε antibody (1,2, 4 µg/mL) of coated plates for 72h. The amount of IL-2 in the supernatant was measured by ELISA and T cell proliferation was measured by CellTrace dilution analyzed by FACS.

### Generation of bone marrow chimera

T cell-depleted bone marrow cells from CD45.2^+^ C57BL/6, Fam49a-KO mice, or Fam49b-KO mice (1x10^6^ cells) were used to reconstitute sublethally irradiated (1000 rad) CD45.1^+^ wild-type mice by i.v. injection. Reconstituted mice were analyzed 8 weeks after bone marrow transfer.

### Cell staining

For cleaved caspase 3 staining [28], homogenized mice thymocyte cells were stained with anti-CCR7/CD197 at a final dilution of 1:50 for 30 min at 37°C prior to additional surface stains. Following surface staining, cells were fixed with Cytofix/Cytoperm (BD Biosciences) for 20 min at 4°C. Cells were then washed with Perm/Wash buffer (BD Biosciences) twice. Cells were stained with anti–cleaved caspase 3 at a 1:50 dilution at 23**°** C for 30 min.

For iNKT staining, Biotinylated PBS-57 loaded or unloaded monomers were obtained from the Tetramer Core Facility of the National Institutes of Health and tetramerized with PE-labeled streptavidin from ProZyme. Hepatic lymphocytes were resuspended in 100 µl of sorter buffer (PBS with 2% FCS, 1 mM EDTA, and 0.1% sodium azide) and stained with PE-iNKT tetramers at a final dilution of 1:200 at 23°C for 30 min.

For transcription factor staining, cells were incubated with surface antibody at 4°C for 20 min, permeabilized at 4°C for 30 min, and then stained with anti–Foxp3 at 23°C for 30 min using a Foxp3/Transcription transcription factor buffer set (Invitrogen, ThermoFisher Scientific). Samples were acquired with BD FACSCelesta (BD Biosciences), and data were analyzed with FlowJo (version 10.7.1).

### Statistical testing

GraphPad Prism was used for all statistical analyses. A nonparametric Mann-Whitney U test or one-way ANOVA was used for estimation of statistical significance. Data is shown as mean ± SEM. *p<0.05, **p<0.01, ***p<0.001, ****p<0.0001.

---

**Supplementary Fig. 1.**
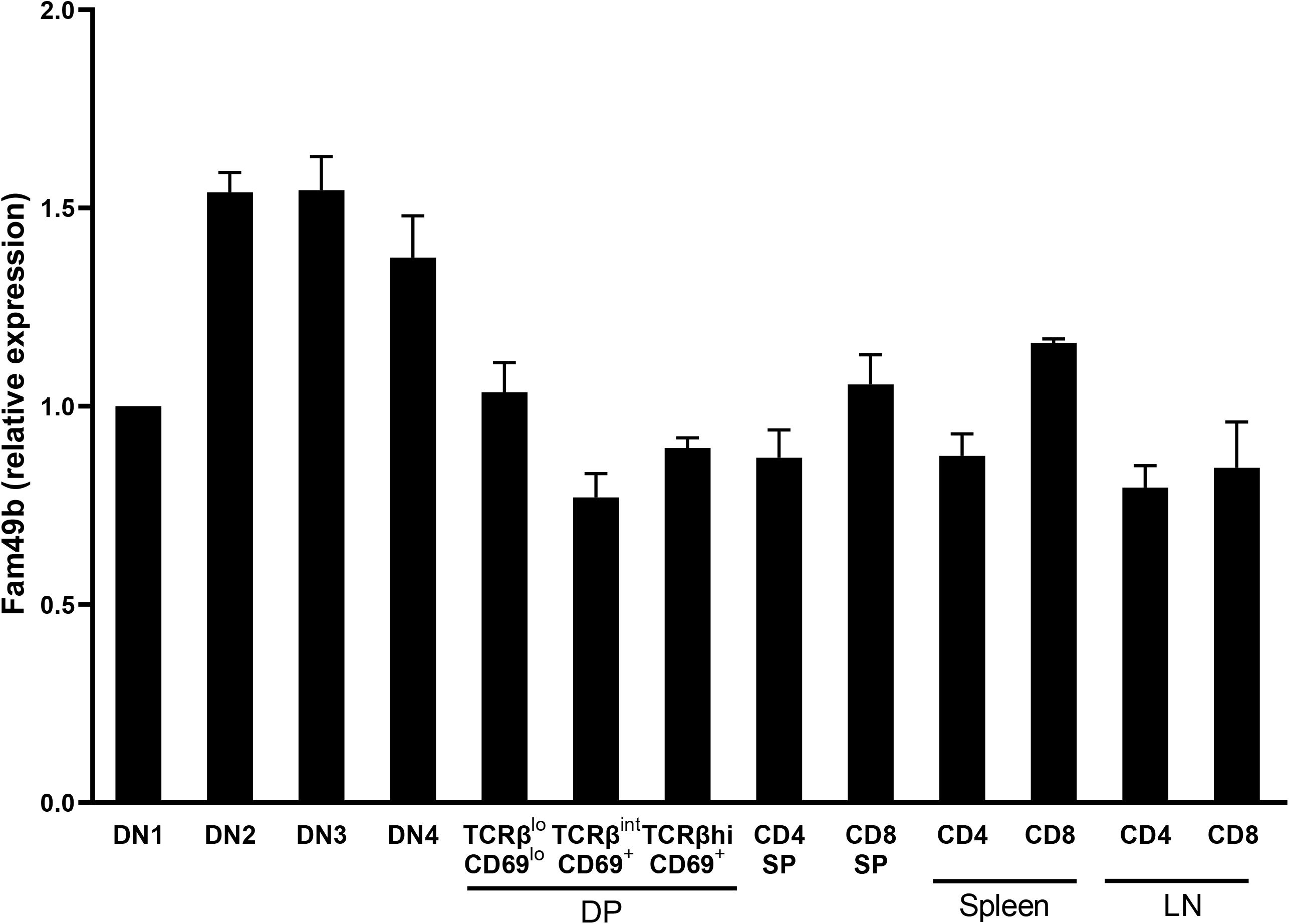
Fam49b expression in thymocyte subsets and T cells from WT mice. (A) Fam49b mRNA expression analyzed by real-time RT-PCR of FACS-sorted subset of WT thymocytes and peripheral T cells. Data shown relative to β actin expression. Error bars denote s.e.m. Data are pooled from two independent experiments. See also Supplementary Figure 1 – source data 6.

**Supplementary Fig. 2.**
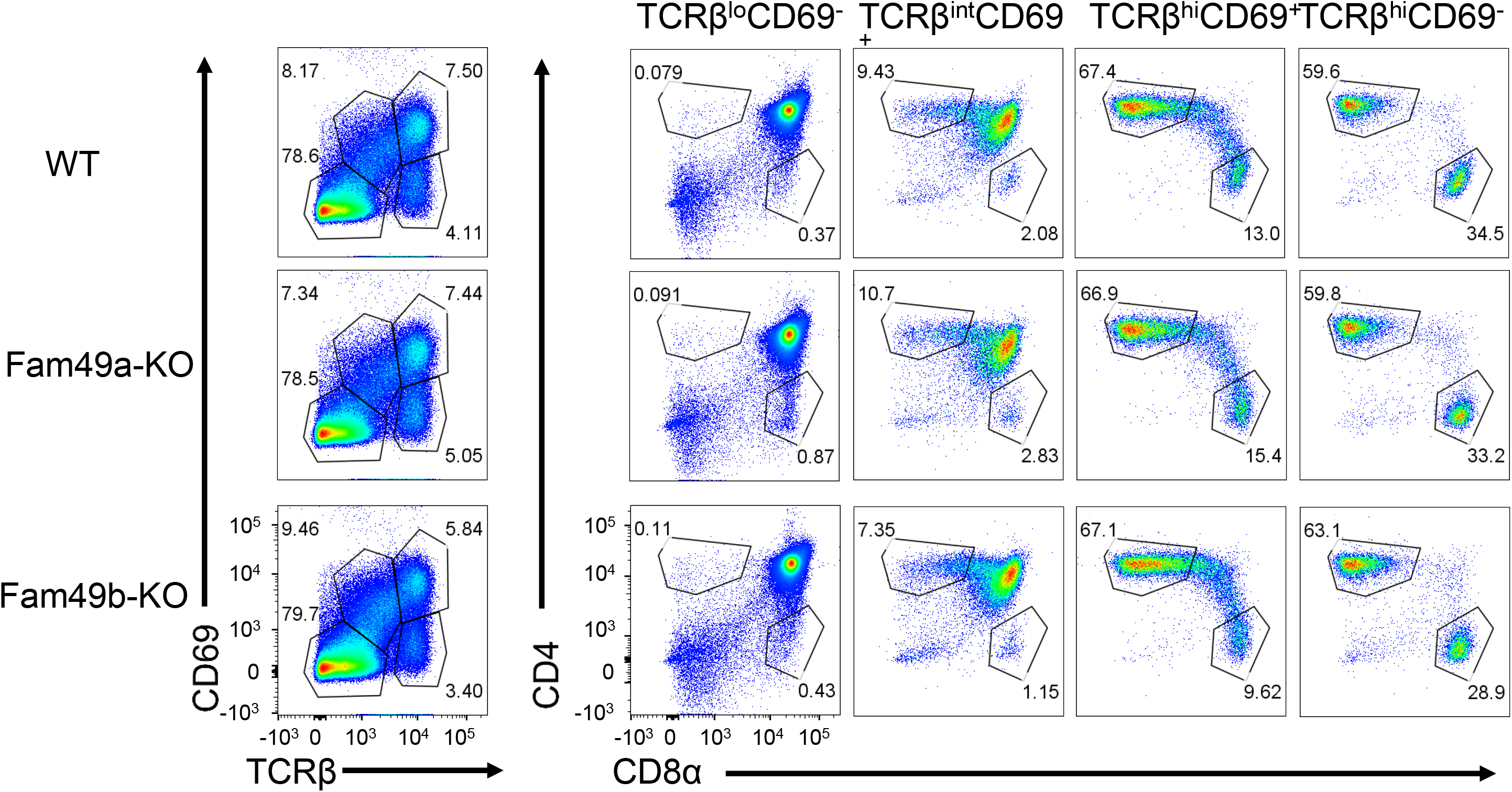
Analyzing thymic selection using TCRβ and CD69 expression in thymus. Representative flow cytometry plot showing TCRβ and CD69 expression (**left**) in total thymocytes from WT, Fam49a-KO, and Fam49b-KO mice. Numbers indicate percentage of CD4 SP or CD8 SP (**right**) from TCRβ by CD69 profile gated (l**eft**). Data are representative of five experiments.

**Supplementary Fig. 3.**
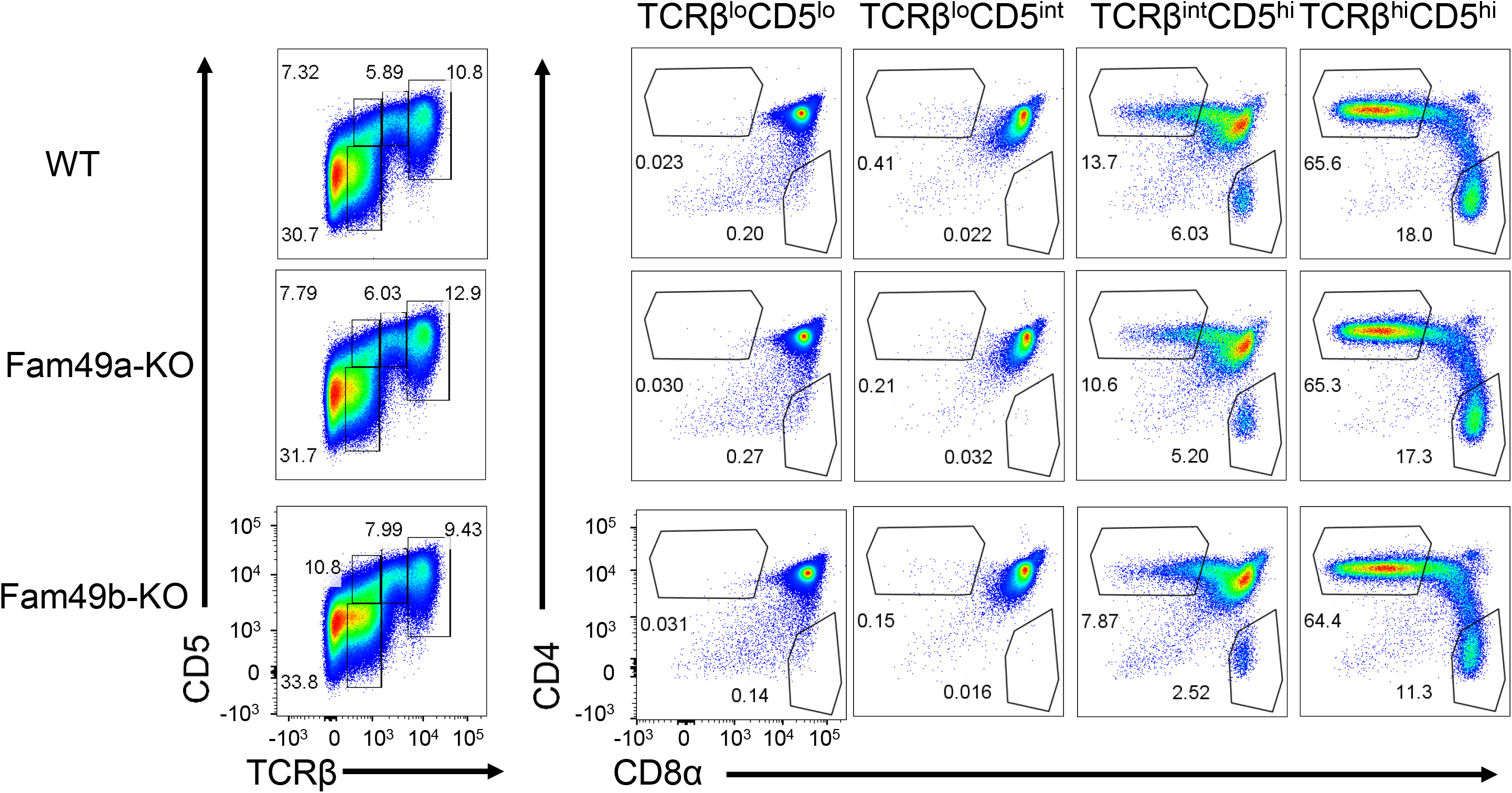
Analyzing thymic selection using TCRβ and CD5 expression in thymus. Representative flow cytometry plot showing TCRβ and CD5 expression (**left**) in total thymocytes from WT, Fam49a-KO, and Fam49b-KO mice. Numbers indicate percentage of CD4 SP or CD8 SP (**right**) from TCRβ by CD5 profile gated (**left**). The Data are representative of five experiments.

**Supplementary Fig. 4.**
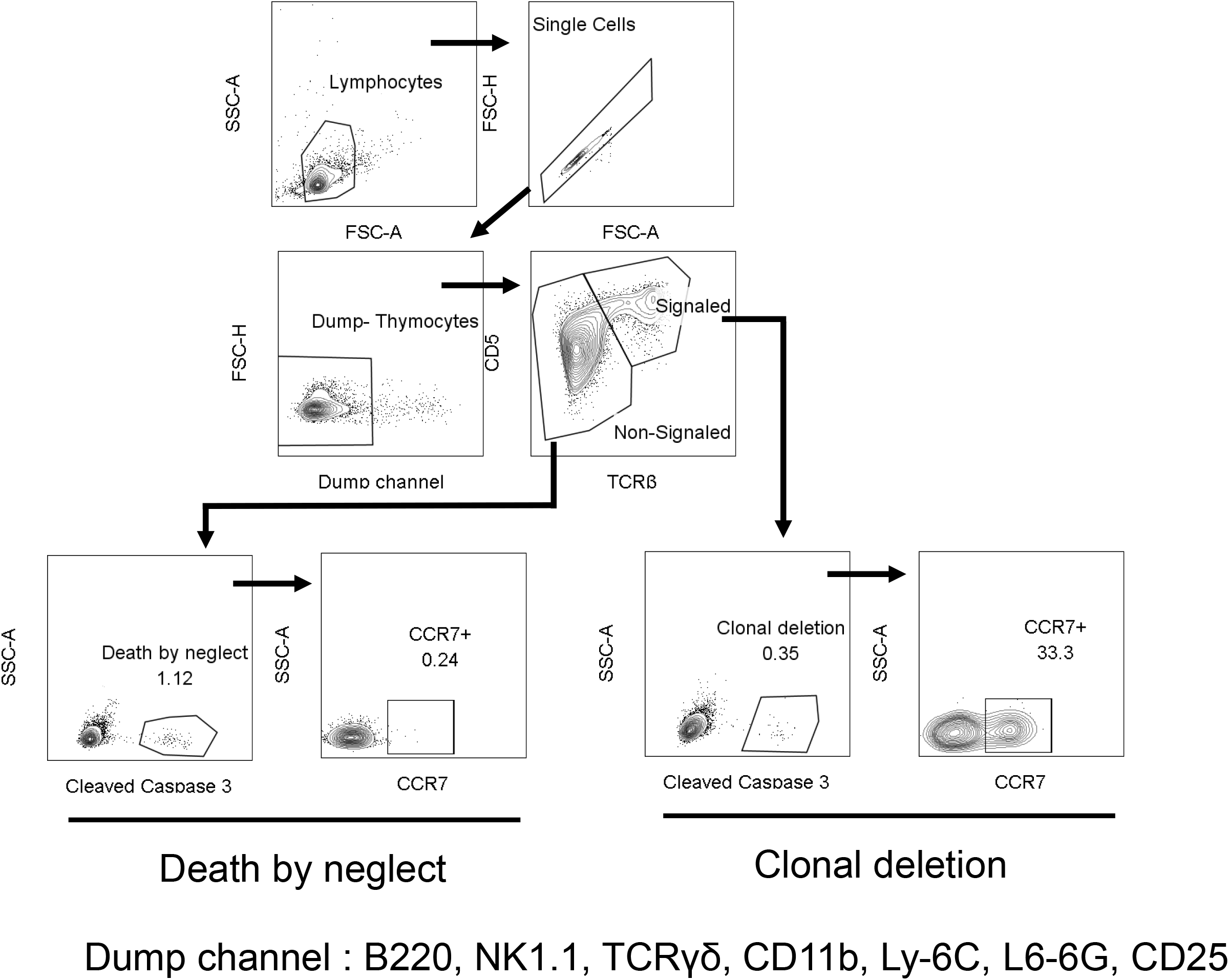
Flow cytometry gating strategies to measure clonal deletion and death by neglect. Signaled and Non-signaled thymocytes identified by TCRβ and CD5 expression, excluding B220^+^, NK1.1^+^, TCRγδ^+^, CD11b^+^, Ly-6C^+^, Ly-6G^+^, CD25^+^ (Dump) cells. Clonal deletion and death by neglect identified by intracellular Cleaved Caspase 3 and anatomic location identified by CCR7. Numbers indicate percentage of cells in each.

**Supplementary Fig. 5.**
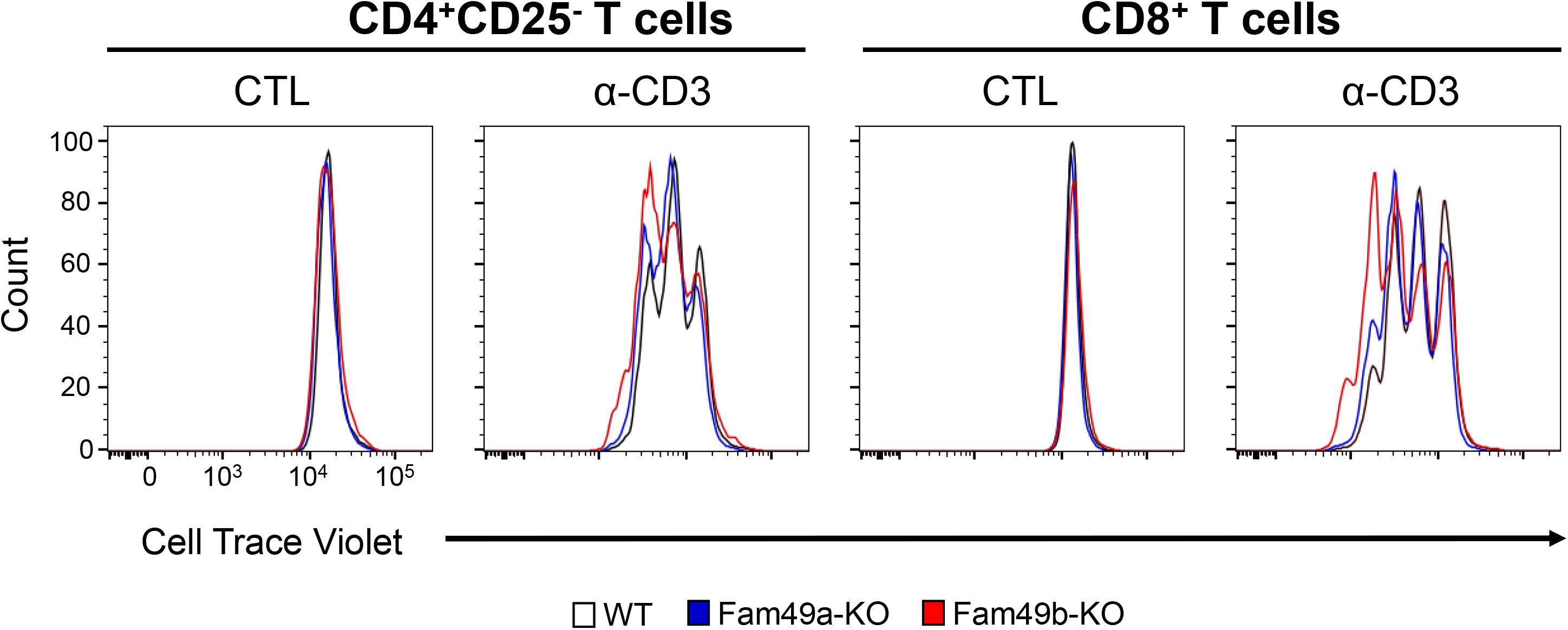
*In vitro* T cell proliferation following α-CD3ε stimulation. Peripheral CD4^+^CD25^-^ T cells (**left**) or CD8^+^ T cells (**right**) were labeled with CellTrace Violet (5 µM, room temperature, 10 min) and stimulated by 4 µg/mL of immobilized anti-CD3ε for 3 days. CellTrace Violet dilution was analyzed by flow cytometry. Data are representative of four experiments.

**Supplementary Fig. 6.**
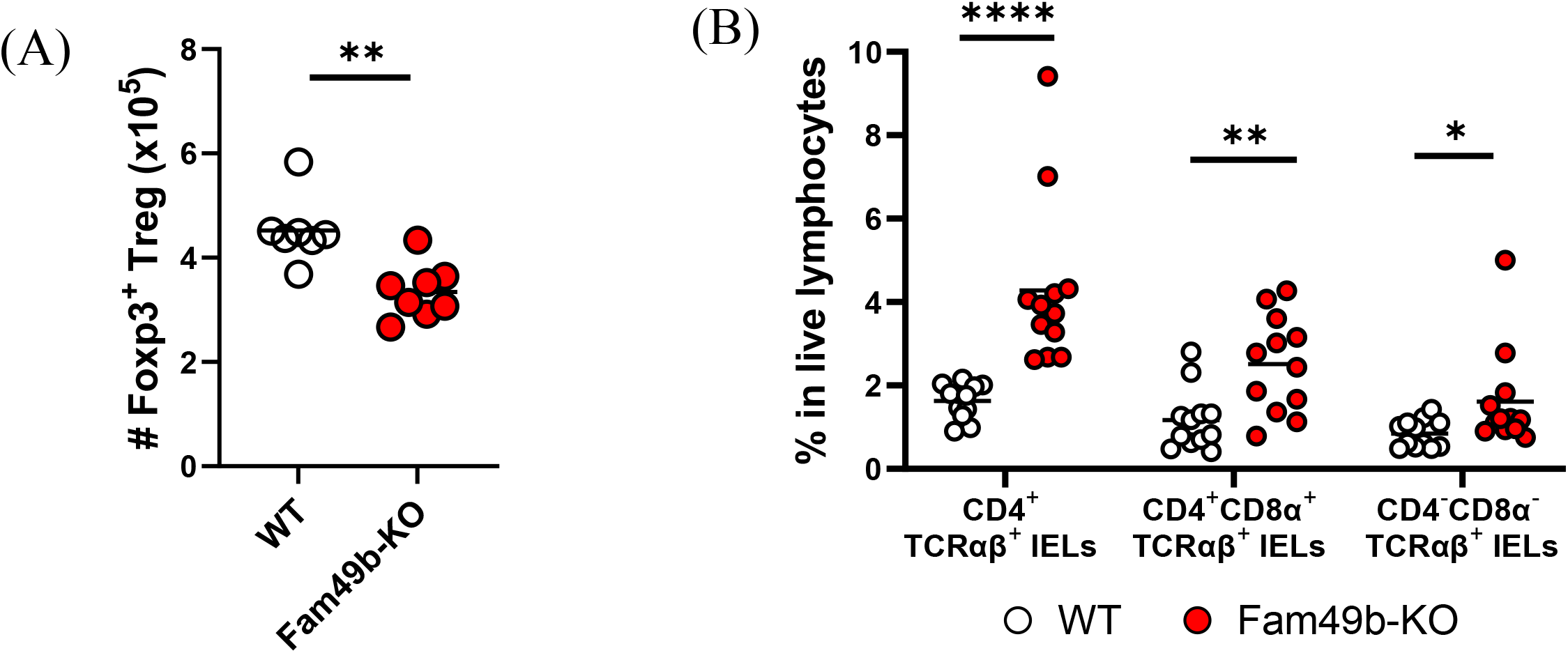
Total number of Treg cells in lymph nodes, and minor IELs T subsets. (A) Total number of Foxp3^+^ regulatory T cells in peripheral lymph nodes from WT and Fam49b-KO mice. Each dot represents an individual mouse. Small horizontal lines indicate the mean of 7-8 mice. **p=0.0012 (Mann-Whitney test). Data are representative from seven independent experiments. See also Supplementary Figure 6 – source data 7. (B) Frequency of CD4^+^TCRαβ^+^ IELs, CD4^+^CD8α^+^TCRαβ IELs, and CD4^-^CD8α^-^TCRαβ IELs T among total live IELs T cells in WT and Fam49b-KO mice. Each dot represents an individual mouse. Small horizontal lines indicate the mean of 12 mice. *p=0.0235 and **p=0.0025 and ****p<0.0001 (Mann-Whitney test). Data are pooled from six independent experiments See also Supplementary Figure 6 – source data 7.

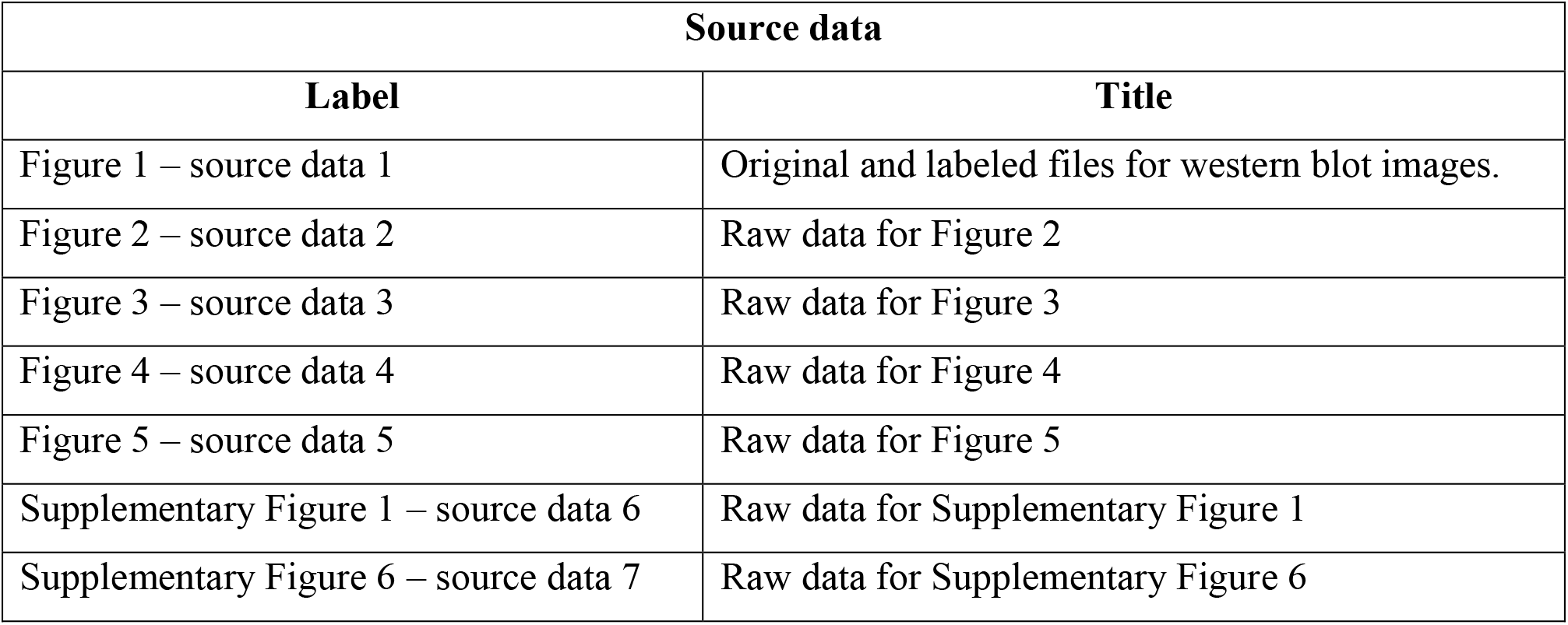

| Source data |  |
| --- | --- |
| Label | Title |
| Figure 1 – source data 1 | Original and labeled files for western blot images. |
| Figure 2 – source data 2 | Raw data for Figure 2 |
| Figure 3 – source data 3 | Raw data for Figure 3 |
| Figure 4 – source data 4 | Raw data for Figure 4 |
| Figure 5 – source data 5 | Raw data for Figure 5 |
| Supplementary Figure 1 – source data 6 | Raw data for Supplementary Figure 1 |
| Supplementary Figure 6 – source data 7 | Raw data for Supplementary Figure 6 |

## Acknowledgments

We thank Dr. J David Peske for editorial comments. Dr. Nilabh Shastri passed away in 2021. May he rest in peace. This work was supported by NIH grant (R01AI130210, R01AI121174, R37AI060040).

## Abbreviations

TCR: T cell receptor

Fam49a: Family with sequence similarity 49 member A

Fam49b: Family with sequence similarity 49 member B

DN: double negative

DP: double positive

SP: single positive

Rho-GTPases: Rho family of small guanosine triphosphatases

GEFs: guanine nucleotide exchange factors

GAPs: GTPase-activating proteins

IELs: intraepithelial lymphocytes

Treg: cells Regulatory T cell

iNKT cells: invariant natural killer T cells

APCs: antigen presenting cells

## References

1. Xu, X., et al., Maturation and emigration of single-positive thymocytes. Clin Dev Immunol, 2013. 2013: p. 282870.

2. Hogquist, K.A., Signal strength in thymic selection and lineage commitment. Curr Opin Immunol, 2001. 13(2): p. 225–31.

3. Gascoigne, N.R., et al., TCR Signal Strength and T Cell Development. Annu Rev Cell Dev Biol, 2016. 32: p. 327–348.

4. Klein, L., et al., Positive and negative selection of the T cell repertoire: what thymocytes see (and don’t see). Nat Rev Immunol, 2014. 14(6): p. 377–91.

5. Hernández-Hoyos, G., et al., Lck activity controls CD4/CD8 T cell lineage commitment. Immunity, 2000. 12(3): p. 313–22.

6. Kappes, D.J., X. He, and X. He, CD4-CD8 lineage commitment: an inside view. Nat Immunol, 2005. 6(8): p. 761–6.

7. Baldwin, T.A., K.A. Hogquist, and S.C. Jameson, The fourth way? Harnessing aggressive tendencies in the thymus. J Immunol, 2004. 173(11): p. 6515–20.

8. Stritesky, G.L., S.C. Jameson, and K.A. Hogquist, Selection of self-reactive T cells in the thymus. Annu Rev Immunol, 2012. 30: p. 95–114.

9. Lambolez, F., M. Kronenberg, and H. Cheroutre, Thymic differentiation of TCR alpha beta(+) CD8 alpha alpha(+) IELs. Immunol Rev, 2007. 215: p. 178–88.

10. Kronenberg, M. and L. Gapin, The unconventional lifestyle of NKT cells. Nat Rev Immunol, 2002. 2(8): p. 557–68.

11. Hsieh, C.S., H.M. Lee, and C.W. Lio, Selection of regulatory T cells in the thymus. Nat Rev Immunol, 2012. 12(3): p. 157–67.

12. Burkhardt, J.K., E. Carrizosa, and M.H. Shaffer, The actin cytoskeleton in T cell activation. Annu Rev Immunol, 2008. 26: p. 233–59.

13. Kumari, S., et al., T cell antigen receptor activation and actin cytoskeleton remodeling. Biochim Biophys Acta, 2014. 1838(2): p. 546–56.

14. Billadeau, D.D., J.C. Nolz, and T.S. Gomez, Regulation of T-cell activation by the cytoskeleton. Nat Rev Immunol, 2007. 7(2): p. 131–43.

15. Kaizuka, Y., et al., Mechanisms for segregating T cell receptor and adhesion molecules during immunological synapse formation in Jurkat T cells. Proc Natl Acad Sci U S A, 2007. 104(51): p. 20296–301.

16. Babich, A., et al., F-actin polymerization and retrograde flow drive sustained PLCγ1 signaling during T cell activation. J Cell Biol, 2012. 197(6): p. 775–87.

17. Babich, A. and J.K. Burkhardt, Coordinate control of cytoskeletal remodeling and calcium mobilization during T-cell activation. Immunol Rev, 2013. 256(1): p. 80–94.

18. Tybulewicz, V.L. and R.B. Henderson, Rho family GTPases and their regulators in lymphocytes. Nat Rev Immunol, 2009. 9(9): p. 630–44.

19. Turner, M., et al., A requirement for the Rho-family GTP exchange factor Vav in positive and negative selection of thymocytes. Immunity, 1997. 7(4): p. 451–60.

20. Fischer, K.D., et al., Defective T-cell receptor signalling and positive selection of Vav-deficient CD4+ CD8+ thymocytes. Nature, 1995. 374(6521): p. 474–7.

21. Zhang, R., et al., Defective signalling through the T- and B-cell antigen receptors in lymphoid cells lacking the vav proto-oncogene. Nature, 1995. 374 (6521): p. 470–3.

22. Dumont, C., et al., Rac GTPases play critical roles in early T-cell development. Blood, 2009. 113(17): p. 3990–8.

23. Guo, F., et al., Rac GTPase isoforms Rac1 and Rac2 play a redundant and crucial role in T-cell development. Blood, 2008. 112(5): p. 1767–75.

24. Phee, H., et al., Pak2 is required for actin cytoskeleton remodeling, TCR signaling, and normal thymocyte development and maturation. Elife, 2014. 3: p. e02270.

25. Shang, W., et al., Genome-wide CRISPR screen identifies FAM49B as a key regulator of actin dynamics and T cell activation. Proc Natl Acad Sci U S A, 2018. 115(17): p. E4051–E4060.

26. Li, R.A., et al., Ndrg1b and fam49ab modulate the PTEN pathway to control T-cell lymphopoiesis in the zebrafish. Blood, 2016. 128(26): p. 3052–3060.

27. Hu, Q., et al., Examination of thymic positive and negative selection by flow cytometry. J Vis Exp, 2012(68).

28. Breed, E.R., M. Watanabe, and K.A. Hogquist, Measuring Thymic Clonal Deletion at the Population Level. J Immunol, 2019. 202(11): p. 3226–3233.

29. McCaughtry, T.M., et al., Clonal deletion of thymocytes can occur in the cortex with no involvement of the medulla. J Exp Med, 2008. 205(11): p. 2575–84.

30. Ueno, T., et al., CCR7 signals are essential for cortex-medulla migration of developing thymocytes. J Exp Med, 2004. 200(4): p. 493–505.

31. Tarakhovsky, A., et al., A role for CD5 in TCR-mediated signal transduction and thymocyte selection. Science, 1995. 269 (5223): p. 535–7.

32. Azzam, H.S., et al., CD5 expression is developmentally regulated by T cell receptor (TCR) signals and TCR avidity. J Exp Med, 1998. 188(12): p. 2301–11.

33. Oh-Hora, M., et al., Agonist-selected T cell development requires strong T cell receptor signaling and store-operated calcium entry. Immunity, 2013. 38(5): p. 881–95.

34. Hogquist, K.A. and S.C. Jameson, The self-obsession of T cells: how TCR signaling thresholds affect fate ‘decisions’ and effector function. Nat Immunol, 2014. 15(9): p. 815–23.

35. Cheroutre, H., F. Lambolez, and D. Mucida, The light and dark sides of intestinal intraepithelial lymphocytes. Nat Rev Immunol, 2011. 11(7): p. 445–56.

36. Gaud, G., R. Lesourne, and P.E. Love, Regulatory mechanisms in T cell receptor signalling. Nat Rev Immunol, 2018. 18(8): p. 485–497.

37. Hwang, J.R., et al., Recent insights of T cell receptor-mediated signaling pathways for T cell activation and development. Exp Mol Med, 2020. 52(5): p. 750–761.

38. Stritesky, G.L., et al., Murine thymic selection quantified using a unique method to capture deleted T cells. Proc Natl Acad Sci U S A, 2013. 110(12): p. 4679–84.

39. McDonald, B.D., et al., Crossreactive αβ T Cell Receptors Are the Predominant Targets of Thymocyte Negative Selection. Immunity, 2015. 43(5): p. 859–69.

40. Fort, L., et al., Fam49/CYRI interacts with Rac1 and locally suppresses protrusions. Nat Cell Biol, 2018. 20(10): p. 1159–1171.

41. Yuki, K.E., et al., CYRI/FAM49B negatively regulates RAC1-driven cytoskeletal remodelling and protects against bacterial infection. Nat Microbiol, 2019. 4(9): p. 1516–1531.

42. Saoudi, A., et al., Rho-GTPases as key regulators of T lymphocyte biology. Small GTPases, 2014. 5.

43. Bosco, E.E., J.C. Mulloy, and Y. Zheng, Rac1 GTPase: a “Rac” of all trades. Cell Mol Life Sci, 2009. 66(3): p. 370–4.

44. Guo, F., et al., Rac GTPase isoforms Rac1 and Rac2 play a redundant and crucial role in T-cell development. Blood, The Journal of the American Society of Hematology, 2008. 112(5): p. 1767–1775.

45. Gomez, M., D. Kioussis, and D.A. Cantrell, The GTPase Rac-1 controls cell fate in the thymus by diverting thymocytes from positive to negative selection. Immunity, 2001. 15(5): p. 703–13.

46. Melichar, H.J., et al., Distinct temporal patterns of T cell receptor signaling during positive versus negative selection in situ. Sci Signal, 2013. 6(297): p. ra92.

47. Pobezinsky, L.A., et al., Clonal deletion and the fate of autoreactive thymocytes that survive negative selection. Nat Immunol, 2012. 13(6): p. 569–78.

48. Ruscher, R., et al., CD8alphaalpha intraepithelial lymphocytes arise from two main thymic precursors . Nat Immunol, 2017. 18(7): p. 771–779.

49. Kurd, N.S., et al., Factors that influence the thymic selection of CD8αα intraepithelial lymphocytes. Mucosal Immunol, 2020.

50. Qiu, Z. and B.S. Sheridan, Isolating Lymphocytes from the Mouse Small Intestinal Immune System. J Vis Exp, 2018(132).

